# Computationally inspired glycoengineering to maximise mAb β4-galactosylation

**DOI:** 10.64898/2026.05.06.723342

**Authors:** Itzcoatl Gomez Aquino, Mina Ghahremanzamaneh, Apostolos Tsopanoglou, Alfonso Blanco, Sara Carillo, Jonathan Bones, Ioscani Jiménez del Val

## Abstract

β4-galactosylation is a critical quality attribute of therapeutic monoclonal antibodies (mAbs), enhancing complement-dependent cytotoxicity, antibody-dependent cytotoxicity, and antibody-dependent cellular phagocytosis. Despite its therapeutic importance, galactosylation remains the most variable glycosylation motif due to its sensitivity to cell culture conditions. Here, we describe a dual genetic engineering strategy applied to two mAb-producing CHO cell lines, DP12 and VRC01, to simultaneously overcome the cellular machinery and metabolic bottlenecks that limit β4-galactosylation. The first engineering event knocks out COSMC, the chaperone required for core 1 β-1,3-galactosyltransferase 1 activity, to redirect UDP-Gal consumption from O-linked β3-galactosylation towards mAb Fc N-linked β4-galactosylation. The second event overexpresses β-1,4-galactosyltransferase 1 (β4GalT1) to augment cellular galactosylation machinery. Each modification was characterised individually (COSMC- and GalT+) and in combination (C-/GT+) across both cell lines in batch and fed batch cultures. The combined C-/GT+ strategy consistently achieved greater than 90% mAb Fc β4-galactosylation, irrespective of host cell line or culture mode. Metabolic characterisation confirmed that both engineering events alleviate their respective bottlenecks: COSMC knockout redirects UDP-Gal flux and β4GalT1 overexpression increases N-galactosylation capacity. The C-/GT+ strategy also reduced production of Man5 glycans, which accelerate serum clearance and pose immunogenicity risks. Metabolic profiling suggests that the COSMC knockout attenuates UTP consumption and contributes to reduced Man5 production. C-/GT+ glycoengineering had no negative impact on mAb titre. Our results establish the C-/GT+ dual glycoengineering strategy as a robust approach for consistently achieving high mAb galactosylation across diverse cell culture conditions, with the additional benefit of reduced Man5 glycans.

**Highlights:**

- Dual COSMC KO and β4GalT1 overexpression achieves >90% mAb Fc galactosylation.
- COSMC KO redirects UDP-Gal from O-glycans to mAb Fc without impacting cell growth.
- Dual glycoengineering reduces production of undesired Man5 glycans.

## 1. Introduction

Monoclonal antibodies (mAbs) represent the most commercially successful class of biopharmaceuticals, with global annual sales exceeding $200 billion since 2021 (Chen et al., 2025; Walsh and Walsh, 2022). Of the more than 200 mAbs currently approved for clinical use, all but one – atezolizumab (Tecentriq^®^), an Fc engineered full-length mAb – carry a conserved N-linked glycan at Asn297 within the CH2 domain of their Fc fragment. Chinese hamster ovary (CHO) cells remain the predominant production host for therapeutic mAbs, accounting for the vast majority of commercial manufacturing processes given to their capacity for human-compatible post-translational modifications, robust growth in suspension culture, and well-established regulatory precedent (Grilo and Mantalaris, 2019).

The Fc N-linked glycan is a critical determinant of mAb therapeutic activity. Glycosylation at Asn297 modulates engagement with Fc gamma receptors (FcγRs) on immune effector cells and with complement component C1q, directly influencing antibody-dependent cell-mediated cytotoxicity (ADCC), antibody-dependent cellular phagocytosis (ADCP), and complement-dependent cytotoxicity (CDC) (van der Horst and Mutis, 2024). Terminal β4-galactosylation is of particular therapeutic relevance, where it enhances CDC activity by facilitating IgG hexamerization at the cell surface, increasing C1q avidity and subsequent complement activation (Peschke et al., 2017; van Osch et al., 2021). mAb Fc β4-galactosylation has been reported to enhance ADCC by up to 30% (Thomann et al., 2016). Recent work has shown that, when combined with afucosylation, increased β4-galactosylation potentiates ADCC through enhanced FcγRIIIA engagement (Zhang et al., 2020). The positive impact of mAb Fc β4-galactosylation on ADCP has also been documented; however, the magnitude of the enhancement varies and may be mAb-dependent (Kuhns et al., 2020).

Despite its therapeutic importance, mAb Fc β4-galactosylation is among the most variable of all Fc glycan features, exhibiting substantial lot-to-lot variability even within the same licensed therapeutic product (Planinc et al., 2017). This variability arises because galactosylation is highly sensitive to cell culture process conditions, including nutrient and trace metal availability, accumulation of metabolic by-products such as ammonium and lactate, dissolved oxygen, pH, and pCO_2_ (Fan et al., 2015; Graham et al., 2019; Hossler et al., 2009; Sumit et al., 2019). Two principal bottlenecks govern the extent of mAb Fc β4-galactosylation in CHO cell culture: (i) the availability of cellular glycosylation machinery, in the form of β-1,4-galactosyltransferase 1 (β4GalT1), and (ii) the availability of uridine diphosphate galactose (UDP-Gal), the nucleotide sugar donor (NSD) that provides the galactose moiety for the reaction. Both bottlenecks have been independently validated: β4GalT1 overexpression in CHO host cells increases mAb Fc β4-galactosylation (Nguyen et al., 2021; Stach et al., 2019), and supplementation of cell culture media with uridine, manganese, and galactose (UMG feeding) expands the intracellular UDP-Gal pool with corresponding improvements in Fc galactosylation (Grainger and James, 2013; Gramer et al., 2011). UDP-Gal transport into the Golgi lumen, mediated by the SLC35A2 NSD transporter, constitutes an additional component of cellular machinery that may further limit galactosylation capacity (Stach et al., 2019).

A crucial but underappreciated constraint on mAb Fc β4-galactosylation is that UDP-Gal is shared with competing cellular glycosylation pathways. Specifically, UDP-Gal is simultaneously consumed by core 1 β-1,3-galactosyltransferase (C1GalT1) to galactosylate cellular O-linked glycans, forming the Core 1 O-glycoform (Galβ1-3GalNAc) that represents the predominant O-glycan structure in CHO cells (Donini et al., 2025; North et al., 2010). Theoretical flux analyses estimate that between 30% and 60% of all UDP-Gal consumed towards glycosylation is directed to cellular O-glycan synthesis (del Val et al., 2016), a figure corroborated by metabolic modelling using CHOmpact – the first metabolic model of CHO cells to explicitly incorporate cellular glycosylation (del Val et al., 2023) – which estimates that approximately 30% of total UDP-Gal flux is consumed by O-galactosylation (unpublished data). This competitive diversion of a shared metabolic resource represents a meaningful and previously unaddressed constraint on mAb Fc β4-galactosylation.

In this study, we address both the cellular machinery and metabolic bottlenecks simultaneously through two orthogonal genetic engineering events applied to mAb-producing CHO-K1 cell lines. The first is knockout of COSMC, the endoplasmic reticulum chaperone specifically required for correct folding and activity of C1GalT1 (Yang et al., 2014). COSMC ablation renders C1GalT1 non-functional, preventing O-linked β3-galactosylation and redirecting the freed UDP-Gal towards mAb Fc N-linked β4-galactosylation. The second engineering event overexpresses β4GalT1 in the production host to overcome the cellular machinery bottleneck. We characterise each modification individually (COSMC- and GalT+) and in combination (C-/GT+), across two mAb-producing CHO-K1 cell lines (DP12 and VRC01) in both batch and fed-batch cultures. The combined C-/GT+ strategy consistently achieves greater than 90% mAb Fc β4-galactosylation irrespective of parental host or culture mode. C-/GT+ engineering also reduces the production of Man5 glycans, which are associated with accelerated serum clearance (Goetze et al., 2011; Yu et al., 2012) and potential immunogenicity risks. Importantly, extended metabolic characterisation suggests that C-/GT+ engineering reduces Man5 production by attenuating UTP consumption and, possibly, Mn^2+^ maldistribution. Together, these findings establish C-/GT+ as a robust, host-agnostic strategy for achieving consistently high mAb Fc β4-galactosylation across diverse manufacturing conditions. The hypergalactosylation achieved by C-/GT+ engineering greatly enhances product consistency and, simultaneously, has the potential to enhance the therapeutic profile of mAbs.

## 2. Material and Methods

### 2.1. Plasmid construction

The plasmid vector pSpCas9(BB)-2A-GFP (hereafter, pX458) was obtained from Addgene (plasmid #48138) (Ran et al., 2013). This plasmid contains a U6 promoter, a scaffold region for a sgRNA cloning site and the coding sequence of Cas9 protein fused with enhanced green fluorescent protein (eGFP). The sgRNA sequence targeting the *COSMC* gene was obtained from Ronda et al. (2014) (listed in the Supplementary Table 1) and was used to generate COSMC- cells. In addition, a plasmid containing the coding sequence of the human β4-galactosyltransferase 1 (hβ4GalT1) was obtained from Invivogen^®^ (Cat. code pUNO1-hb4galt1). In this construct, expression of hβ4GalT1 is driven by a human elongation factor promoter (hEF1). For successful isolation of hβ4GalT1+ cells, pUNO1-hb4galt1 contains a blasticidin resistance cassette (blasticidin-S deaminase, *BsrS2*), the expression of which is driven by a human cytomegalovirus (hCMV) promoter. The sequencing primers for pUNO, pUNO1-hb4galt1, and the hβ4GalT1 gene are presented in Supplementary Tables 2 and 3.

### 2.2. Cell Culture and transfection

Two mAb-producing cell lines were used to evaluate the glycoengineering strategies in this study: CHO-DP12 clone #1934 (ATCC^®^ CRL-12445™), producing an anti-interleukin-8 mAb (Heinrich et al., 2011) and CHO-VRC01 that expresses a broadly neutralising antibody (bNAb) directed at HIV-1 CD4 binding sites (Wu et al., 2010). Both antibodies are humanised IgG1 molecules and contain a conserved glycosylation site at the Asn297 residue of their Fc fragment. The VRC01 mAb also contains an additional glycosylation site on the Fab fragment (Reddy et al., 2025). CHO DP12 was purchased from LGC Standards (Middlesex, UK) and was adapted to serum-free suspension culture at the National Institute for Bioprocessing Research and Training (NIBRT). CHO VRC01 was kindly provided by the Vaccine Research Centre of the US National Institutes of Health. CHO-DP12 and CHO-VRC01 were routinely passaged in Ex-Cell® 302 media (Merck, Cat. No. 14324C) and ActiPro media (HyClone, Cat. No. SH31039.02), respectively. The media were supplemented with 4 mM of L-glutamine (Merck, Cat. No. K0283-BC) and 200 or 100 nM of methotrexate (Sigma-Aldrich, Cat. No. 454126) for DP12 and VRC01, respectively. Cells were propagated in static mode at 37 °C and 5% CO_2_ in a Steri-Cycle CO_2_ incubator (Thermo Scientific, Waltham, US), with a working volume of 5-10 mL, at an initial viable cell density (VCD) of 0.2 million cells/mL and until a VCD between 1 and 1.5 million cells/mL was reached. Cells were subcultured every 2-3 days for a minimum of three passages before transfection or characterisation. VCD and viability were determined through dye exclusion with a haematocytometer and 0.4% (w/v) trypan blue (Gibco, Life Technologies, 15250061).

Transfections were carried out in an Amaxa^®^ CLB-Transfection device (Lonza, Basel, CH), using an Amaxa^®^ CLB-Transfection Kit (Lonza, Cat. No. VECA-1001). The transfection protocol followed manufacturer instructions. Briefly, 1 to 5 million cells/mL growing in exponential phase were centrifuged at 500×g for 10 min and resuspended in the transfection solution along with 3-5 μg of purified endotoxin-free plasmid, the mixture was placed in the supplied cuvette and pulsed in the device (program ‘Cell line 9’). Immediately after transfection, 0.5 mL of the appropriate pre-warmed media was added to the cuvette, and the entire content was recovered and transferred into 6-well plates containing 1.5 mL of preconditioned cell culture media. The cells were incubated for 24 h, after which, the media was replaced. Four days post-transfection, cells transfected with pX458 and its derivative plasmids were evaluated for eGFP expression using an Olympus CKX inverted fluorescence microscope (Olympus Corporation, Tokyo, JP) equipped with a mercury/xenon burner and a 510/50 nm filter. Cells were then bulk sorted using a FACSAria III™ or a FACSMelody™ cell sorter (Becton-Dickinson, Franklin Lakes, US) equipped with a 488 nm excitation laser and a 510/50 nm filter. On day 4 post-transfection, cells modified with the pUNO1-hb4galt1 plasmid were subjected to an antibiotic selection protocol, where cells were exposed to blasticidin-supplemented media (10 μg/mL). During this process, cells were centrifuged at 500×g for 10 min and resuspended in fresh media every 48 to 72 hours. The selection process concluded when VCD reached 0.5-1.0 million cells/mL and viability was greater than 95%.

### 2.3. Cell culture characterisation

Cell culture characterisation experiments were performed in batch and fed-batch mode in 125 mL Erlenmeyer shake flasks with vented caps and a 30 mL working volume. Flasks were cultured in a Steri-Cycle CO_2_ incubator (Thermo Scientific, Waltham, US) at 5% CO_2_, 37 °C, and shaken at 120 rpm (Thermo Fisher Scientific, Cat. No. 88881102). All cultures were seeded at 0.2 million cells/mL and were terminated when viability fell below 70%. 1 mL samples were taken daily to determine VCD, viability, metabolite profiles, and mAb titre.

For the fed-batch cultivations, feeding schedules and recipes were established according to vendor instructions. CHO-DP12 cells cultured in Ex-Cell^®^ 302 (Merck, Cat. No. 14324C) were supplemented with 6.25% (v/v) of Ex-Cell^®^ Advanced™ CHO Feed 1 (Merck, Cat. No. 24367C) using the starting cell culture volume (30 mL) as reference and were fed every 48 hours starting on day 3 post-seeding. CHO-VRC01 cells were cultured in HyClone^TM^ ActiPro^TM^ media (Cytiva, SH31039), supplemented daily from day 3 with 3.0% (v/v) HyClone^TM^ Cell Boost^TM^ 7a (Cytiva, SH31026) and 0.3% (v/v) HyClone^TM^ Cell Boost^TM^ 7b (Cytiva, SH31027). For both cell lines, a 45% (w/v) glucose solution was fed into the culture aiming to maintain a residual glucose concentration above 4.0 g/L (>22 mM). Daily in-process glucose measurements were made using an Accu-Chek^®^ Aviva glucose meter (Roche Diagnostics, Manheim, DE) and test strips (Roche Diagnostics, Cat. No. 0643970). Residual concentrations of glucose, lactate, glutamine, glutamate, and ammonia were measured with a Cedex Bio Analyser (Roche Diagnostics, Manheim, DE). mAb titre was determined with Protein A HPLC, using a mAbPac^TM^ column (Thermo Scientific, Cat. No. 082539) fitted to an Agilent 1200 HPLC (Agilent Technologies, Santa Clara, US).

On the final day of culture, ∼15mL were harvested by spinning cells down at 3,000×g for 10 minutes and purifying the supernatant on an ÄKTA Start^TM^ FPLC (Cytiva, Uppsala, SE). mAb was captured with a HiTrap^®^ protein A column (Cytiva, 17040201) and, after elution, pH was neutralised with 200μL of 1M Tris-HCL (pH 9). Purified mAb concentration was determined with a ND-100 Nanodrop^TM^ spectrophotometer (Thermo Fisher Scientific, Paisley, UK).

Fc-glycosylation of the purified mAb samples was analysed using the single-chain Fc glycoprofiling method developed by Carillo et al. (2020). Briefly, mAb samples were digested with the immunoglobulin-degrading enzyme of *Streptococcus pyogenes* (IdeS). The digested subunits were then run on a Vanquish^TM^ Flex binary UHPLC equipped with a MAbPac RP column (2.1 x 50 mm, 4 µm particle size, Thermo Fisher Scientific, Sunnyvale, US) and coupled to a Q Exactive^TM^ Plus Hybrid Quadrupole Orbitrap mass spectrometer (Thermo Fisher Scientific, Bremen, DE) for high resolution accurate mass acquisition of Fc subunits. Raw data were deconvoluted using Xtract algorithm available on BioPharma Finder software (Thermo Fisher Scientific, San Jose, US).

### 2.4. HPLC analysis of nucleotides and nucleotide sugar donors (NSDs)

The cell pellet resulting from nutrient/metabolite sampling was resuspended in ice-cold 50% v/v acetonitrile:water, incubated on ice for 10 min, and then spun down for 5 minutes at 18,000×g and 4°C. The supernatant was stored at -80°C before further processing. After thawing, samples were dried with a SpeedVac vacuum concentrator (Savant, Thermo Fisher Scientific, Paisley, UK), resuspended in 2 mL of 10 mM NH_4_HCO_3_ (Acros, Cat. No. 393210010), and subjected to ion-pairing solid-phase extraction (IP-SPE) using Supelclean™ ENVI-Carb™ columns (Sigma-Aldrich, 57088). The IP-SPE columns were conditioned with 3 mL of 80% acetonitrile (ACROS, Cat. No. 258560025) in 0.1% trifluoroacetic acid (Alfa Aesar, Cat. No. L06374 AP), followed by 2 mL of water. Samples were loaded onto the columns and washed with 2 mL of water, 2 mL of 25% acetonitrile, 200 μL of water, and 2 mL of 50 mM triethylamine acetate buffer, pH 7 (Sigma-Aldrich, Cat. No. 90358). 2 mL of 25% acetonitrile in 50 mM triethylamine acetate buffer (pH 7.0) were used to elute nucleotides and NSDs. Eluates were lyophilised and stored at -80°C prior the HPLC analysis.

Nucleotide and NSDs were quantified using the ion-pairing reverse phase HPLC method described by Nakajima et al. (2010) on an Agilent 110 series HPLC system (Agilent Technologies, Santa Clara, US). Buffer A was 100 mM potassium phosphate buffer supplemented with 8 mM of the tetrabutylammonium hydrogen sulphate (J.T Baker, Cat No. 15598904) ion-pairing reagent. Buffer B was a 70% v/v Buffer A with 30% v/v acetonitrile. Separation was performed at 40°C on an Inertsil ODS-4 column (particle size = 3 μml 10×4.6 mm internal diameter; GL Sciences, Tokyo, JP). The cellular extracts were dissolved in 200μL of deionised water and injected to the column equilibrated with Buffer A. A constant flow rate of 1 mL/min was maintained and detection was performed at 254 nm. The gradient elution is described in Supplementary Table 4. For quantification, calibration curves and spiked samples were run with 21 nucleotides and nucleotide sugars (Supplementary Tables 5 and 6).

### 2.5. Lectin-aided flow cytometry

Lectin-aided flow cytometry was employed to identify the cell population with the COSMC knock-out. The lectin used to detect the COSMC- phenotype was Vicia Villosa (VVL, Vector Labs, B-1235), which has a unique affinity towards exposed N-acetylglucosamine of truncated O-glycans (the so-called ‘Tn antigen’). The lectin was conjugated with fluorescein isothiocyanate (FITC) which has excitation/emission wavelengths of 495/519 nm. For lectin staining, 3-5 million cells were pelleted, resuspended in 500 μL PBS (Thermo Fisher Scientific, Cat. No. 10010023), and incubated with 1 μL VVL-FITC for 20 min on ice and in the dark. After incubation, cells were washed by spinning them down at 1,800×g for 4 mins and resuspending them in 500 μL of PBS (Thermo Fisher Scientific, Cat. No. 10010023). Stained cells were bulk sorted with a FACSAria III™ or a FACSMelody™ cell sorter (Becton-Dickinson, Franklin Lakes, US), and the recovered pool was propagated and banked.

Changes in cell surface N-linked β4-galactosylation (CSG) were also determined using lectin-aided flow cytometry, using a protocol adapted from Gramer et al. (2011). Here, a 3-5 million cell pellet was resuspended in 500 μL of 1M HEPES buffer (Sigma-Aldrich, H0887) and incubated with 15 μg / 10^6^ cells Texas Red labelled *Erythrina cristagalli* lectin (ECL) (ECL-TR; EyLabs, T-5901-2) at 37°C for 20 mins in the dark. After incubation, three washing steps were performed, where cells were spun down at 1,800×g for 4 mins and resuspended in 500 μL of 1M HEPES buffer (Sigma-Aldrich, H0887). Stained samples were analysed on a CytoFLEX LX flow cytometer (Beckman Coulter, La Brea, US) using a yellow laser and a 610/20 filter. Data from 10,000 singlet events were acquired and recorded.

### 2.6. Statistical analysis and calculations

Statistical analysis was performed using GraphPad Prism 11.0.0. The level of significance was set at p < 0.05 using ANOVA followed by Tukey’s test for pairwise mean comparisons. *, **, ***, and **** denote p ≤ 0.05, p ≤ 0.01, p ≤ 0.001, and p ≤ 0.0001, respectively. To compare CSG of ECL-TR stained populations, the staining index (SI) was computed using Equation 1, where MFI(+) and MFI(-) are the median fluorescence intensities of the stained and unstained samples, respectively and rSD is the standard deviation of the unstained sample.

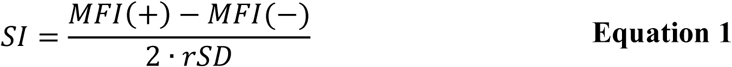

% galactosylated species were calculated with Equation 2, where A2G1F denotes a monogalactosylated and fucosylated biantennary glycan, A2G2F represents the bi-galactosylated and fucosylated biantennary glycan, and A2S1G1F represents a biantennary fucosylated glycan with terminal galactose on one antenna and terminal sialic acid on the other. % non-galactosylated was calculated with Equation 3, where Man5 is five mannose containing glycan and A2G0F is the fucosylated biantennary glycan with terminal N-acetylglucosamine.

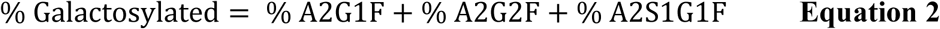

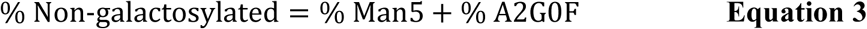

## 3. Results

### 3.1. Generation of CHO DP12 and CHO VRC01 glycovariant cell pools

The cell engineering strategy proceeded as follows: first, parental cells were subjected to CRISPR/Cas9-mediated knockout of the Core 1 β3-galactosyltransferase-specific chaperone (COSMC) using the p458 system (Ran et al., 2013), in which Cas9 is fused to GFP, to generate COSMC- variants. GFP-positive cells were initially enriched by flow cytometry, after which cells lacking GFP signal were selected by FACS to minimise stable Cas9 integration. A COSMC knockout phenotype was further enriched using lectin-assisted flow cytometry with *Vicia villosa* lectin, which specifically binds the Tn antigen (the non-galactosylated Core 1 O-glycan). Screening data for COSMC knockout enrichment are shown in Supplementary Figures 1 (DP12) and 2 (VRC01).

Parental cells were transfected with the pUNO plasmid encoding human β4-galactosyltransferase 1 (hβ4GalT1) and selected with blasticidin to generate GalT+ variants. COSMC- cells were transfected with the same pUNO-hβ4GalT1 construct and selected with blasticidin to produce dual-engineered C-/GT+ cells. Following selection, integration of the hβ4GalT1 sequence was confirmed by genomic DNA extraction and PCR amplification using gene-specific primers, with results shown in Supplementary Figures 3 (DP12) and 4 (VRC01). Collectively, these approaches enabled the successful generation of stable COSMC-, GalT+, and C-/GT+ cell pools for both DP12 and VRC01 mAb-producing CHO cell lineages.

### 3.2. C-/GT+ glycoengineering drives increased mAb β4-galactosylation in batch culture

The glycoengineered cells derived from CHO DP12 and CHO VRC01 cells were cultured in batch mode. The glycoengineered variants cultured were: Parental (non-engineered cells), COSMC- (cells where the Core 1 β3-galactosyltransferease-specific molecular chaperone has been knocked out using CRISPR/Cas9), GalT+ (cells where human β4-galactosyltransferase 1 has been ectopically expressed), and C-/GT+ (cells that have been modified with simultaneous COSMC knockout and ectopic hβ4GalT1 expression). Shake flask cultures were performed, and samples were taken to monitor VCD, mAb titre, and endpoint mAb Fc glycoprofiling.

In batch cultures, no significant differences were observed in glycosylation profiles produced by Parental and COSMC- CHO DP12 cells (Figure 1A). A sharp increase in β4-galactosylated glycans was observed in GalT+ and C-/GT+ cells, reaching 77.9% ± 1.5% A2G2F species in GalT+ cells and 78.6% ± 2.8% in C-/GT+ cells (Figure 1A). Interestingly, a modest yet statistically significant difference (p = 0.041) is observed in the fraction of A2G0F species produced by GalT+ (7.5% ± 0.3%) and C-/GalT+ (4.0% ± 0.6%) cells. Although statistically significant differences in individual β4-galactosylated glycans were not observed, the reduced production of A2G0F by C-/GT+ cells resulted in significantly higher overall galactosylation (96.0% ± 0.6%) when compared with GalT+ cells (92.5% ± 0.3%) (Figure 1B).

**Figure 1.**
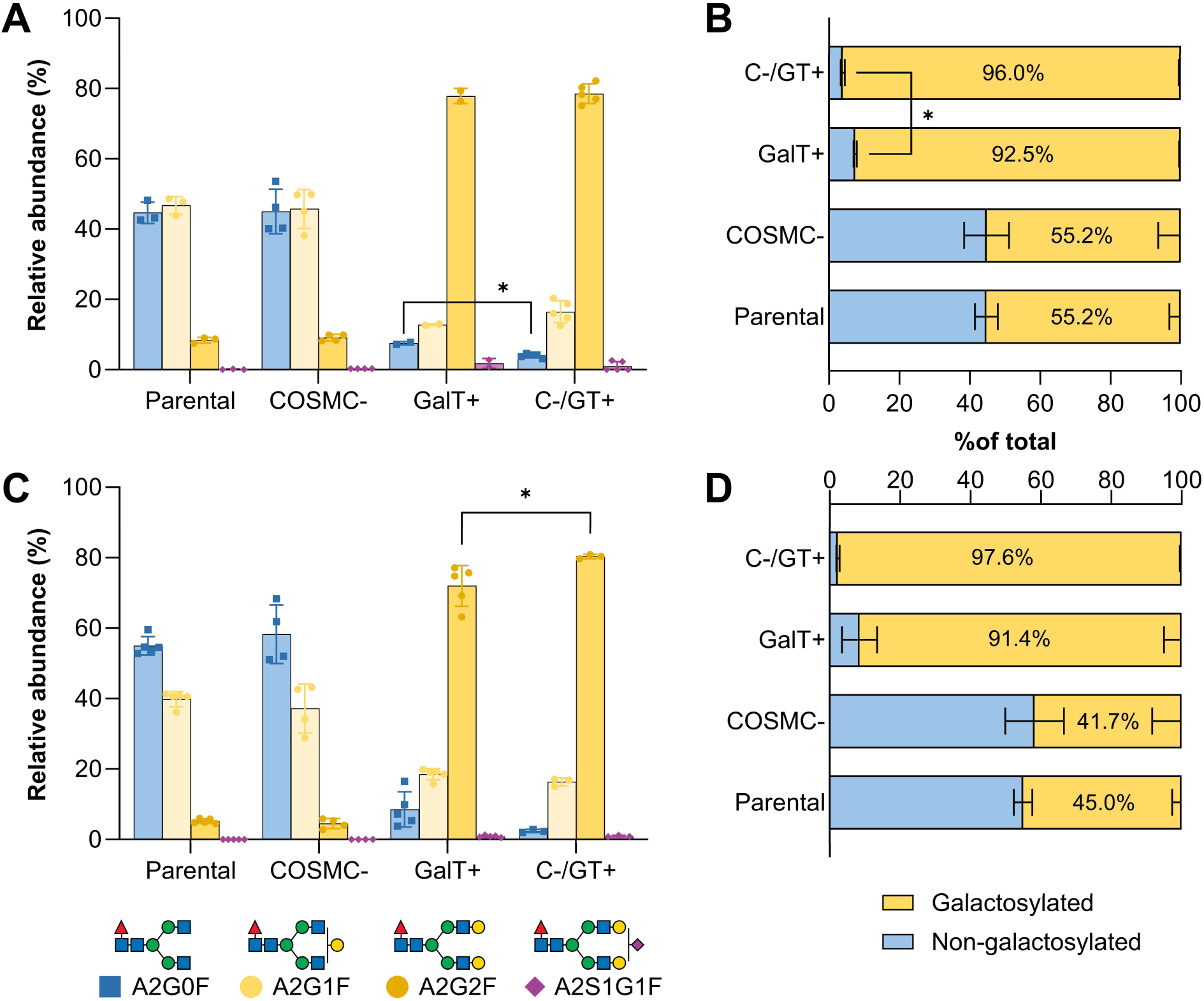
mAb Fc glycosylation profiles produced by parental and glycoengineered DP12 and VRC01 cells cultured in batch mode. mAb Fc glycoforms from batch cultures of Parental, COSMC-, GalT+, and C-/GT+ DP12 (A) and VRC01 (C) are shown, with total galactosylated species summarised in (B) and (D). In (A) and (C), points represent individual biological replicates (relative Fc glycan abundance) and bars indicate means; glycan identities follow CFG nomenclature (Varki et al., 2015). In (B) and (D), yellow bars denote total galactosylated species (A2G1F + A2G2F + A2S1G2F), and blue bars indicate non-galactosylated A2G0F. Two-way ANOVA with Tukey’s post hoc test (GraphPad Prism 11.0.0) was performed to assess mean differences between groups. Significance: *p < 0.05, **p < 0.01, ***p < 0.001, ****p < 0.0001.

Batch cultures of CHO VRC01 cell variants produced similar results as CHO DP12. No significant differences were observed among the mAb Fc glycoforms produced by Parental and COSMC- cells Figure 1C. A considerable increase in mAb Fc β4-galactosylation is also observed in VRC01 the GalT+ and C-/GT+ variants, where β4GalT1 is ectopically expressed Figure 1C. A statistically significant difference (p = 0.011) is observed in the A2G2F mAb Fc glycoform produced by GalT+ (72.0% ± 4.6%) and C-/GT+ (80.3% ± 0.6%) cells. No difference is observed in overall galactosylated glycoforms (Figure 1D), likely due to the high variability in A2G0F and A2G2F glycans produced by VRC01 GalT+ cells.

These results suggest a scheme where β4GalT availability initially limits β4-galactosylation. Once the β4GalT bottleneck is overcome through ectopic expression, UDP-Gal availability becomes limiting. This is consistent with the results presented in Figure 1. No differences in β4-galactosylation are observed between Parental and COSMC- cells, and it is only when β4GalT is ectopically expressed that a substantial increase in mAb Fc β4-galactosylation is observed (Parental and COSMC- vs. GalT+ in Figure 1). The further increase in β4-galactosylation produced by cells containing both modifications (C-/GT+) would correspond to additional UDP-Gal being made available for mAb β4-galactosylation through ablation of cellular O-linked β3-galactosylation. Overall, the batch culture glycoprofiling results suggest that the dual C-/GT+ glycoengineering strategy maximises mAb Fc β4-galactosylation by simultaneously alleviating cellular machinery (β4GalT)and metabolic (UDP-Gal) bottlenecks.

### 3.3. Glycovariant performance in batch cultures

Small differences are observed in the VCD profiles across CHO DP12 glycoengineered variants. Growth of Parental and COSMC- cells is indistinguishable up to 144 h of culture, after which a drop in Parental cell VCD is observed (Figure 3A). GalT+ and C-/GT+ cells present slightly higher growth rates during the first 96 h of culture, with C-/GT+ cells presenting a higher peak cell density (4.23 ± 0.31 10^6^ cells/mL) and integral of viable cell density (495.6 ± 39.7 10^6^ cells h/mL) (Figure 2A and Table 1), suggesting improved cell health.

**Figure 2.**
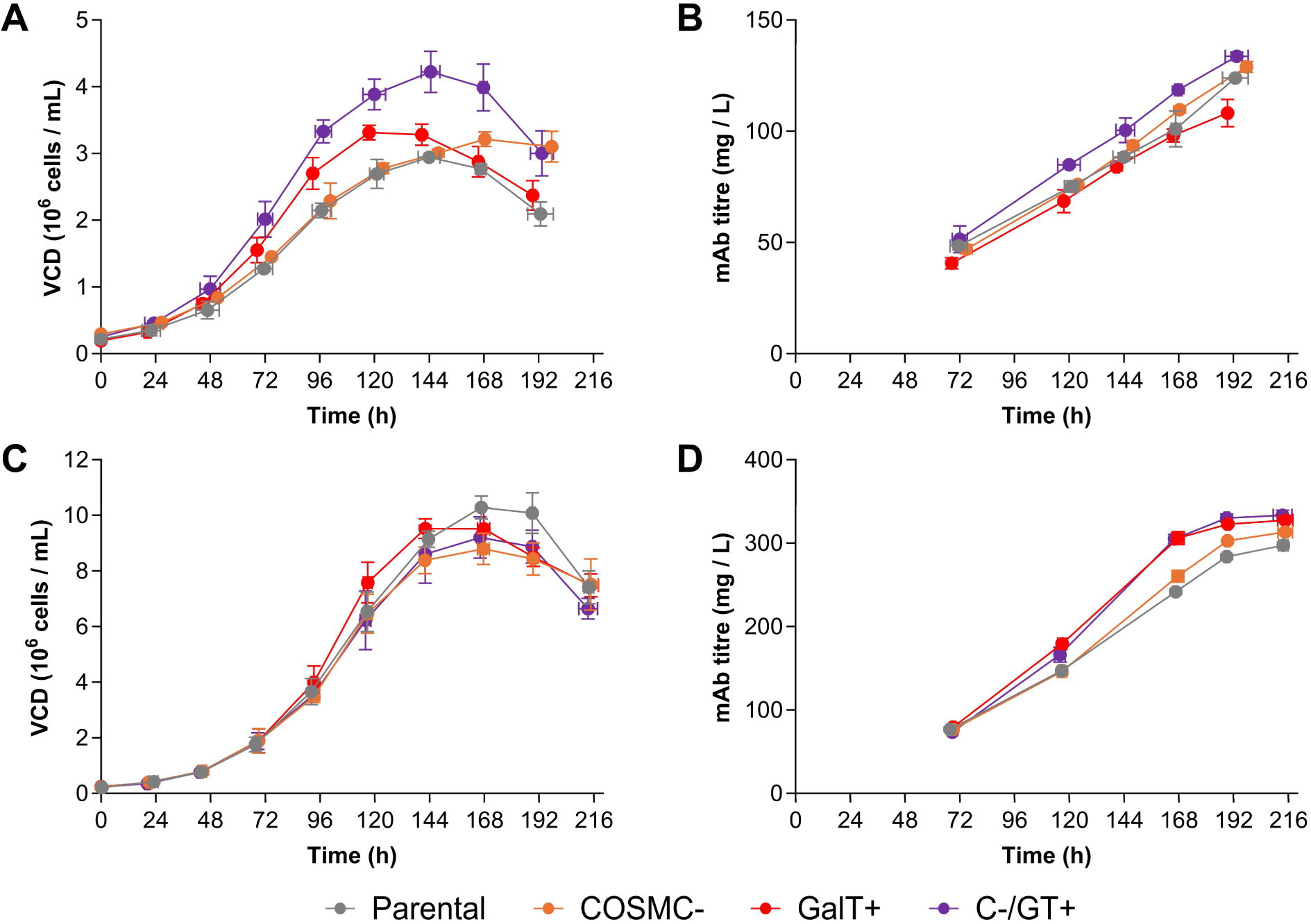
VCD and Titre in batch cultures of glycoengineered DP12 and VRC01 cells. Viable cell density (VCD) profiles for batch cultures of Parental (grey), COSMC- (orange), GalT+ (red), and C-/GT+ (purple) CHO DP12 (A) and CHO VRC01 (C) are shown. Corresponding mAb titre profiles for CHO DP12 (B) and CHO VRC01 (D) are also presented.

**Figure 3.**
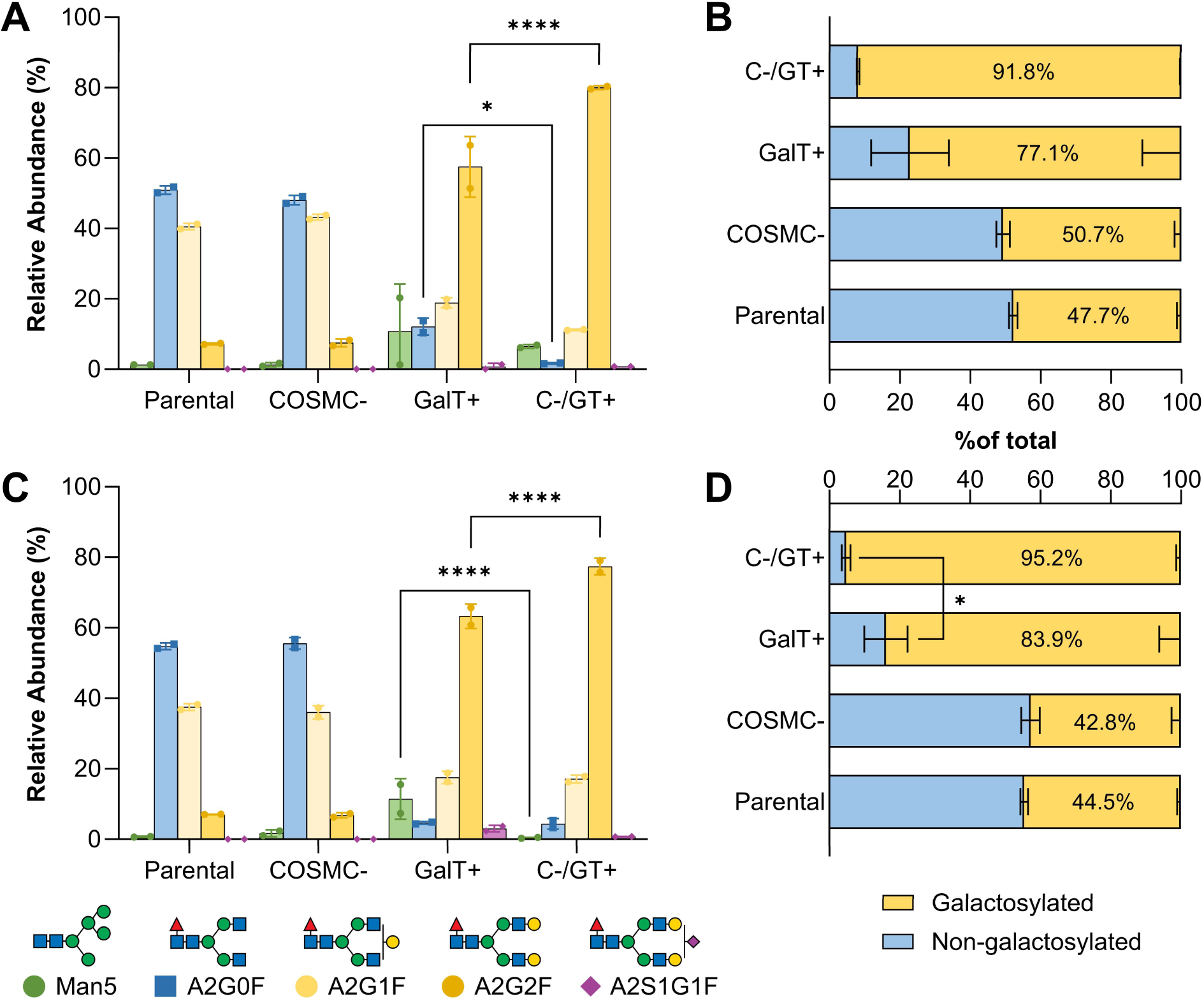
mAb Fc glycosylation profiles produced by parental and glycoengineered CHO DP12 and CHO VRC01 cells cultured in fed batch mode. mAb Fc glycoforms produced in fed batch cultures of Parental, COSMC-, GalT+, and C-/GT+ variants of CHO DP12 (A) and CHO VRC01 (C) are shown. Total galactosylated glycoforms are summarized in (B) and (D). In (A) and (C), points represent the relative abundance of each detected Fc glycan from individual cultures, and bars indicate mean values ± S.D. of biological duplicates. Glycan identities are annotated using CFG nomenclature (Varki et al., 2015). In (B) and (D), yellow bars denote mean total galactosylated species (A2G1F + A2G2F + A2S1G2F), while blue bars represent non-galactosylated glycans (Man5 + A2G0F). GraphPad Prism 11.0.0 was used to perform two-way ANOVA with Tukey’s post hoc test. Significance is indicated as *p < 0.05, **p < 0.01, ***p < 0.001, ****p < 0.0001.

**Table 1.**
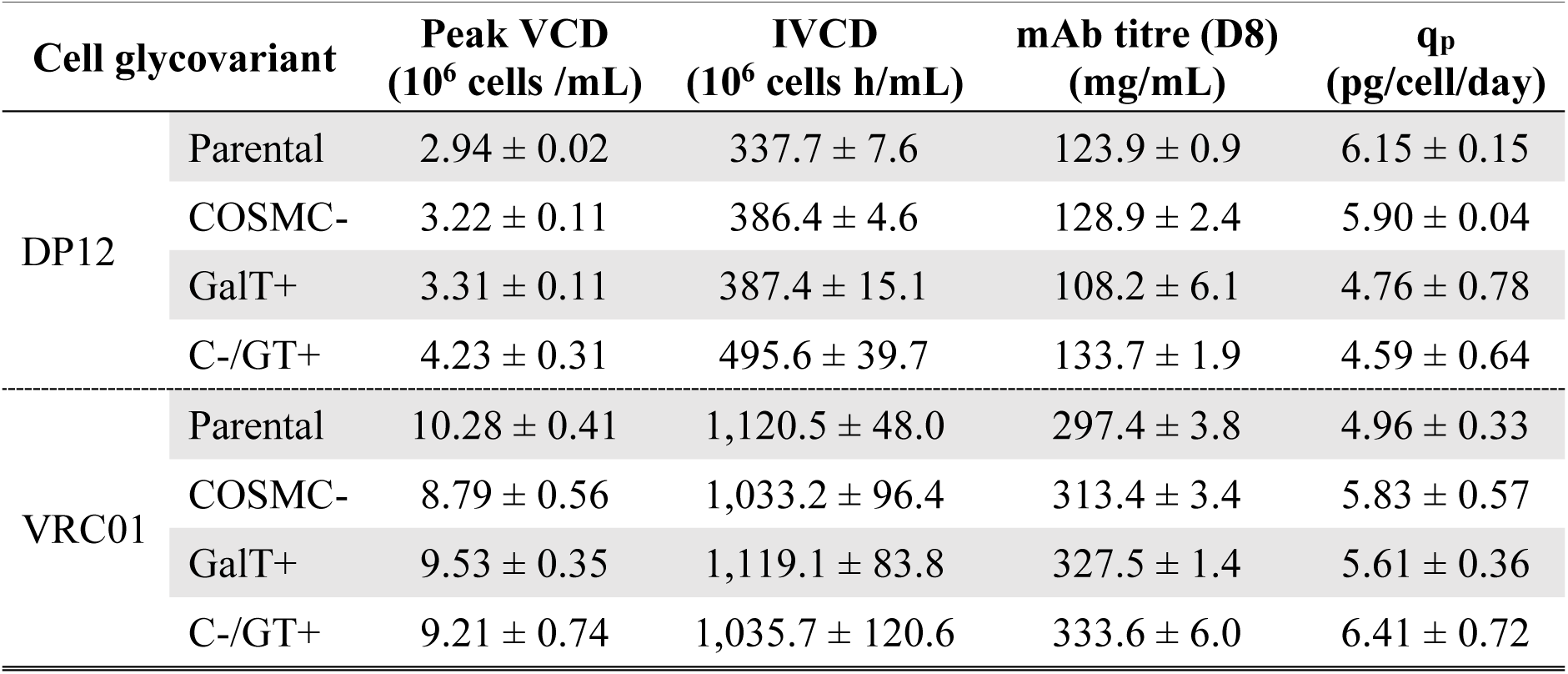
Batch culture IVCD, mAb titre, and q_p_ of CHO DP12 and VRC01 glycovariants.

| Cell glycovariant | | Peak VCD<br>($10^6$ cells /mL) | IVCD<br>($10^6$ cells h/mL) | mAb titre (D8)<br>(mg/mL) | $q_p$<br>(pg/cell/day) |
| --- | --- | --- | --- | --- | --- |
| DP12 | Parental | $2.94 \pm 0.02$ | $337.7 \pm 7.6$ | $123.9 \pm 0.9$ | $6.15 \pm 0.15$ |
| | COSMC- | $3.22 \pm 0.11$ | $386.4 \pm 4.6$ | $128.9 \pm 2.4$ | $5.90 \pm 0.04$ |
| | GalT+ | $3.31 \pm 0.11$ | $387.4 \pm 15.1$ | $108.2 \pm 6.1$ | $4.76 \pm 0.78$ |
| | C-/GT+ | $4.23 \pm 0.31$ | $495.6 \pm 39.7$ | $133.7 \pm 1.9$ | $4.59 \pm 0.64$ |
| VRC01 | Parental | $10.28 \pm 0.41$ | $1,120.5 \pm 48.0$ | $297.4 \pm 3.8$ | $4.96 \pm 0.33$ |
| | COSMC- | $8.79 \pm 0.56$ | $1,033.2 \pm 96.4$ | $313.4 \pm 3.4$ | $5.83 \pm 0.57$ |
| | GalT+ | $9.53 \pm 0.35$ | $1,119.1 \pm 83.8$ | $327.5 \pm 1.4$ | $5.61 \pm 0.36$ |
| | C-/GT+ | $9.21 \pm 0.74$ | $1,035.7 \pm 120.6$ | $333.6 \pm 6.0$ | $6.41 \pm 0.72$ |

Minor differences in mAb titre at harvest were observed across the glycoengineered variants of CHO DP12 cells. All glycovariants achieved equivalent or marginally higher mAb titre at harvest with the exception of GalT+, where a shortfall of 12.65% ± 0.72% was observed (Figure 2B and Table 1). A reduction in cell specific productivity (q_p_) was observed in the DP12 glycovariant cells (Table 1), which may be associated with the VCD profiles and inversely correlates with peak cell density (Table 1). Accelerated growth by GalT+ and C-/GT+ cells leads to lower q_p_ (4.76 ± 0.78 pcd and 4.59 ± 0.64 pcd, respectively), while the lower growth rate by Parental and COSMC- cells achieves higher q_p_ (6.15 ± 0.15 pcd and 5.90 ± 0.04 pcd, respectively) (Table 1). Cell specific productivity is known to correlate inversely with growth rate in CHO cells due to rebalancing of cellular resources from product to cell biosynthesis (Gutierrez et al., 2020).

CHO VRC01 glycovariants present more consistent VCD profiles than CHO DP12. Growth up to 144 h of culture is nearly identical for all four VRC01 glycovariants, after which Parental cells continue to grow while the VCD for COSMC-, GalT+, and C-/GT+ begins to decline (Figure 2C). No significant differences are observed in peak cell density or IVCD across all VRC01 glycovariants (Table 1). Only minor differences in endpoint mAb titre are observed, where COSMC-, GalT+, and C-/GT+ produce 5.39% ± 0.08%, 10.13% ± 0.14%, and 12.18% ± 0.27% more mAb than Parental cells, respectively (Figure 2D and Table 1). These variations in mAb titre and consistent IVCDs translate into increased q_p_ for the glycoengineered variants of CHO VRC01 cells (Table 1). While differences in titre and q_p_ may reflect normal cell culture variability, the time profiles in Figure 2D indicate that the GalT+ and C-/GT+ VRC01 variants behave differently from Parental and COSMC- cells.

### 3.4. C-/GT+ glycoengineering increases mAb β4-galactosylation in fed batch culture

To further test if the knockout of COSMC contributes to increasing UDP-Gal availability towards N-linked β4-galactosylation, a fed batch cultures were performed with DP12 and VRC01 glycovariants. Our reasoning was that the cells would be more metabolically challenged during prolonged culture periods. No significant difference is observed between Parental and COSMC- variants in both DP12 and VRC01 cells (Figure 3A and C). As in batch cultures, a considerable increase in β4-galactosylation is observed in hβ4GalT1-expressing cells (Figure 3B and Figure 3D). Notably, a sizable and variable amount of Man5 is produced by DP12 and VRC01 GalT+ cells, whereas Man5 was not detected in batch cultures.

The most interesting comparison is between GalT+ and C-/GT+ cells because it demonstrates how the COSMC knockout contributes to enhancing mAb β4-galactosylation across longer culture periods where the cells may be more metabolically challenged. In contrast to batch cultures, larger differences are observed between DP12 GalT+ and C-/GT+ cells for the A0G0F and A2G2F glycans. DP12 GalT+ cells produce 10.4% ± 1.7% more non-galactosylated A0G0F (p = 0.046) and 22.5% ± 6.1% less bi-galactosylated A2G2F (p < 0.0001) than C-/GT+ cells (Figure 3A). VRC01 cells present a similar trend, where GalT+ cells yield 11.0% ± 4.1% more Man5 (p < 0.0001) and 14.1% ± 3.0% less A2G2F (p < 0.0001) than C-/GT+ cells (Figure 3C). These differences result in 11.3% ± 2.8% lower overall mAb Fc β4-galactosylation (p = 0.044) (Figure 3D). Together, our glycoprofiling results suggest that, when combined with ectopic hβ4GalT1 expression, knocking out COSMC maximizes mAb β4-galactosylation.

### 3.5. Glycovariant performance in fed batch cultures

VCD and mAb titre profiles for DP12 and VRC01 glycovariants are presented in Figure 4, and values for peak VCD, IVCD, and q_p_ for these cultures are shown in Table 2. In DP12 cells, the VCD profiles of Parental and COSMC variants are similar and reach a higher peak values and IVCD than the hβ4GalT1-expressing GalT+ and C-/GT+ variants (Figure 4A and Table 2). The mAb titre profiles of DP12 variants contrast with VCD. The lowest titre is observed in GalT+ variants, while the highest occurs in COSMC- cells. Parental and C-/GT+ cells present very similar profiles (Figure 4B). The contrast between VCD and mAb titre profiles results in q_p_ differences across DP12 glycovariants, where COSMC- achieves the highest q_p_ and GalT+ cells the lowest (Table 2). Notably, q_p_ is, on average, 60% lower and IVCD is 2.4-fold higher in fed batch cultures of DP12 variants, indicating considerable re-routing of metabolic resources towards cell growth at the expense of productivity.

**Figure 4.**
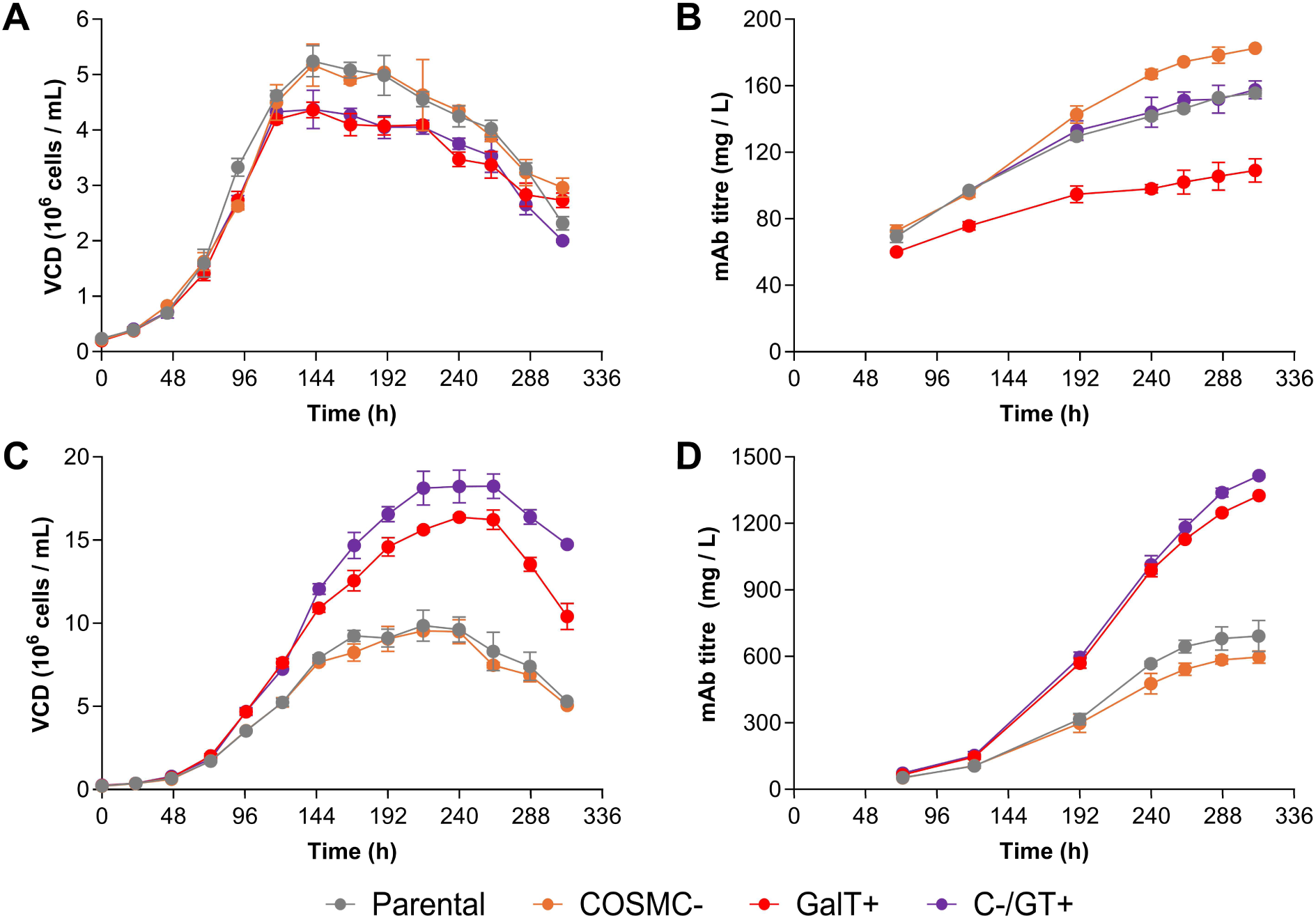
VCD and Titre in fed batch cultures of glycoengineered DP12 and VRC01 cells. Viable cell density (VCD) profiles for batch cultures of Parental (grey), COSMC- (orange), GalT+ (red), and C-/GT+ (purple) CHO DP12 (A) and CHO VRC01 (C) are shown. Corresponding mAb titre profiles for CHO DP12 (B) and CHO VRC01 (D) are also presented.

**Table 2.** Fed batch culture IVCD, mAb titre, and q_p_ of CHO DP12 and VRC01 glycovariants.

| Cell glycovariant | | Peak VCD<br>($10^6$ cells/mL) | IVCD<br>($10^6$ cells h/mL) | mAb titre (D8)<br>(mg/mL) | $q_p$<br>(pg/cell/day) |
| --- | --- | --- | --- | --- | --- |
| DP12 | Parental | $5.24 \pm 0.28$ | $1,045.4 \pm 41.0$ | $155.6 \pm 2.1$ | $2.07 \pm 0.04$ |
| | COSMC- | $5.17 \pm 0.38$ | $1,031.2 \pm 11.9$ | $182.6 \pm 0.9$ | $2.69 \pm 0.04$ |
| | GalT+ | $4.36 \pm 0.14$ | $897.1 \pm 13.9$ | $109.0 \pm 7.0$ | $1.37 \pm 0.15$ |
| | C-/GT+ | $4.37 \pm 0.35$ | $905.5 \pm 3.6$ | $157.6 \pm 5.4$ | $2.47 \pm 0.07$ |
| VRC01 | Parental | $9.86 \pm 0.93$ | $1,807.7 \pm 106.9$ | $691.4 \pm 70.7$ | $8.71 \pm 0.43$ |
| | COSMC- | $9.54 \pm 0.08$ | $1,730.8 \pm 51.2$ | $596.7 \pm 27.5$ | $7.78 \pm 0.16$ |
| | GalT+ | $16.38 \pm 0.14$ | $2,881.3 \pm 19.5$ | $1,325.7 \pm 1.4$ | $10.71 \pm 0.06$ |
| | C-/GT+ | $18.24 \pm 0.74$ | $3,268.8 \pm 73.6$ | $1,415.7 \pm 7.2$ | $10.03 \pm 0.20$ |

Profiles for glucose, lactate, glutamine, glutamate, and ammonia for DP12 fed batch cultures are presented in Supplementary Figure 5. Parental and COSMC- cells show higher glucose consumption due to their increased VCD. No differences are observed in lactate profiles across all DP12 variants, suggesting that Parental and COSMC- cells have more efficient central carbon metabolism. All four DP12 variants present similar ammonia profiles which, when coupled with the lower glutamate accumulation of Parental and COSMC- cells, suggests that these variants may have lower glutamine anaplerosis towards TCA. Overall, Parental and COSMC- DP12 cells appear to have more efficient central carbon metabolism that translates into higher proliferation and lower q_p_. Regardless of the differences observed across DP12 variants, the dual C-/GT+ glycoengineering strategy produces mAb titres that are equivalent to those achieved by Parental cells, indicating no shortfall in overall culture performance.

The effects of cell glycoengineering on growth, productivity, and metabolism are considerably different and more marked in VRC01 cells. GalT+ and C-/GT+ cells present substantially higher VCD and IVCD than Parental and COSMC- cells (Figure 4C and Table 2). Interestingly, GalT+ and C-/GT+ VRC01 variants achieve a two-fold higher mAb titre than Parental and COSMC- cells, reaching above 1.3 g/L (Figure 4D and Table 2). The increase in mAb titre is not only due to higher IVCD: GalT+ and C-/GT+ cells present a ∼20% higher q_p_. Metabolic profiles for VRC01 variants are presented in Supplementary Figure 6, where GalT+ and C-/GT+ cells present very similar behaviour. Both variants present lower residual glucose concentrations and produce lower lactate levels than Parental and COSMC- cells. hβ4GalT expressing cells also show increased glutamine accumulation, lower glutamate profiles, and equivalent ammonia profiles than the Parental and COSMC- variants. These profiles indicate that GalT+ and C-/GT+ cells exhibit considerably more efficient metabolism that, in turn, translates into enhanced growth and increased mAb productivity.

Although enhanced culture performance arises in cell pools that have undergone blasticidin resistance selection, it is unlikely that presence of the blasticidin resistance (BsdR) cassette itself underlies the effect – cell pools generated with the pUNO vector lacking the hβ4GalT gene had similar batch culture performance as Parental and COSMC- cells (Supplementary Figure 7). The increased performance of hβ4GalT-expressing cells is therefore more likely due to pleiotropic effects of the selection process, and/or broader host-cell consequences of β4GalT overexpression on cellular glycosylation. β4GalT overexpression likely alters cellular glycosylation, which in turn, may affect cellular behaviour (Narimatsu et al., 2021).

### 3.6. Glycoengineering impacts cell surface β4-galactosylation

To assess the effect of the glycoengineering strategies on cellular proteins, cell surface N-linked β4-galactosylation was quantified with flow cytometry using Texas Red-labelled *Erythrina cristagalli* lectin (ECL), which specifically binds to β4-linked galactose residues. Figure 5A presents day 6 CSG data for DP12 variants cultured in batch mode, while Figure 5C shows day 6 CSG data for batch cultured VRC01 variants.

**Figure 5.**
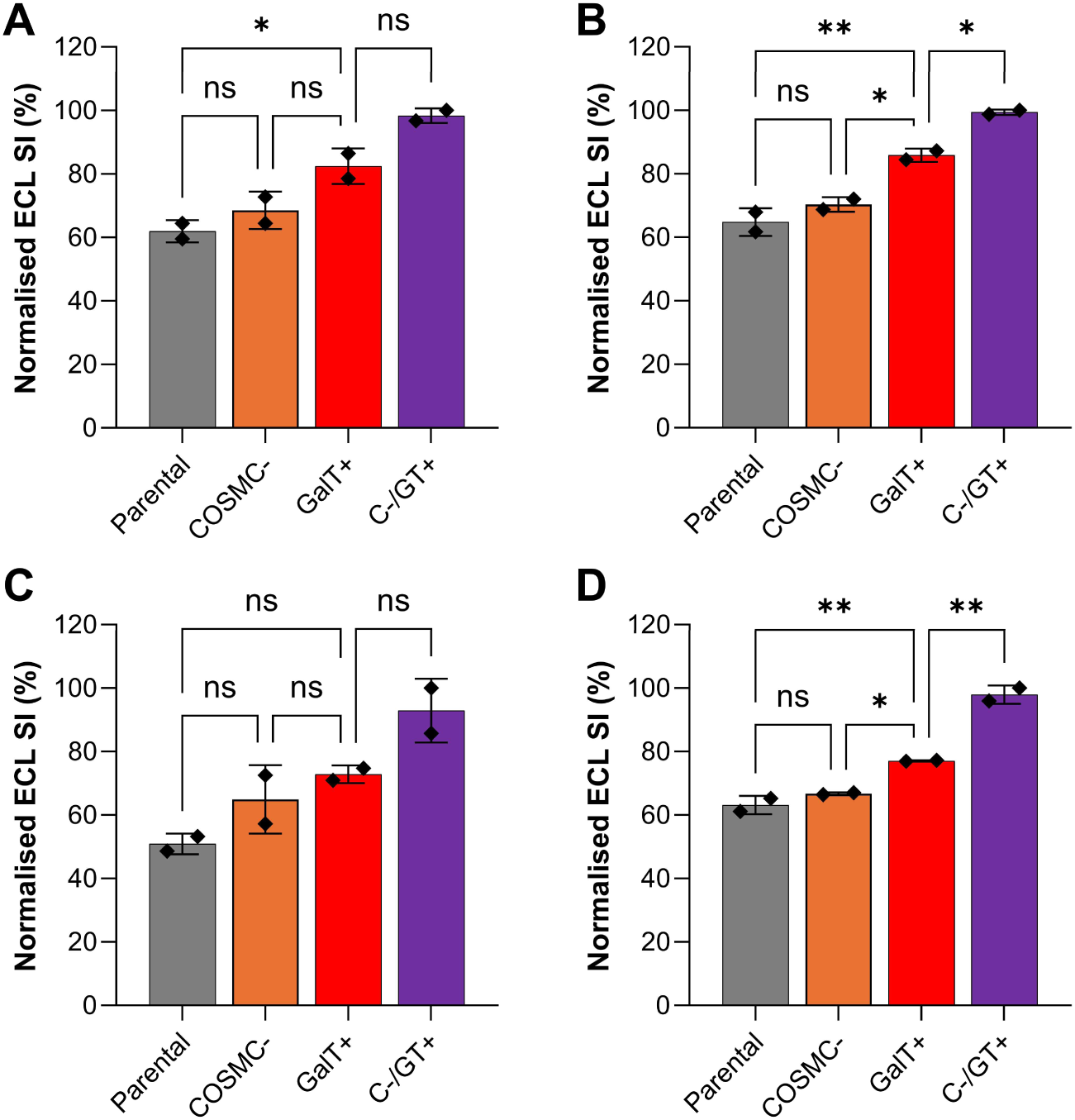
Cell surface β4-galactosylation for DP12 and VRC01 glycovariants cultured under batch and fed batch mode. Normalised staining index (SI) of DP12 glycovariants in batch (A) and fed-batch (B), and VRC01 glycovariants in batch (C) and fed-batch (D). Samples were collected on day 6 (batch) or day 12 (fed-batch), stained with Texas Red-labelled *Erythrina cristagalli* lectin (ECL), and analysed with flow cytometry. Data are mean ± S.D. (N = 2) and were normalised to the maximum SI within each group. A one-way ANOVA with Tukey’s post hoc test was performed (GraphPad Prism 11.0.0). Significance: *p < 0.05, **p < 0.01, ***p < 0.001, ****p < 0.0001.

Figure 5B and Figure 5D present day 12 CSG data for fed batch cultures of DP12 and VRC01 variants, respectively. No statistically significant differences are observed between Parental and COSMC- variants, which is consistent with the mAb Fc glycoprofiling results shown in Figure 1 and Figure 3. A CSG increase of 20.5% ± 6.6% (p = 0.036) is observed between Parental and GalT+ DP12 variants (Figure 5A), while no significant difference was found in batch cultured VRC01 (p = 0.142), likely due to variability associated with low non-normalised SI values obtained (Supplementary Figure 8). Although omitted from Figures 5A and 5C for clarity, batch cultures of DP12 and VRC01 variants showed significantly higher CSG in C-/GT+ variants compared to parental cells (DP12: p = 0.005; VRC01: p = 0.019).

Fed batch cultures followed similar increasing CSG trends (Parental < GalT+ < C-/GT+) as those observed in batch cultures (Figures 6B and 6D); however, higher statistical significance was found, which was likely due to lower variability. As in batch cultures, no differences were found between fed batch cultured Parental and COSMC- variants (Figures 6B and 6D). Between Parental and GalT+ variants, CSG increased significantly in DP12 (p = 0.005) and VRC01 (p = 0.009) cells. An additional increase in CSG was observed between fed batch cultures of GalT+ and C-/GT+ variants of DP12 (p = 0.026) and VRC01 (p = 0.002) (Figures 5B and 5D). These CSG results show that the glycoengineering events impact cellular glycosylation and may, possibly, underly differences found in cell growth and metabolism.

**Figure 6.**
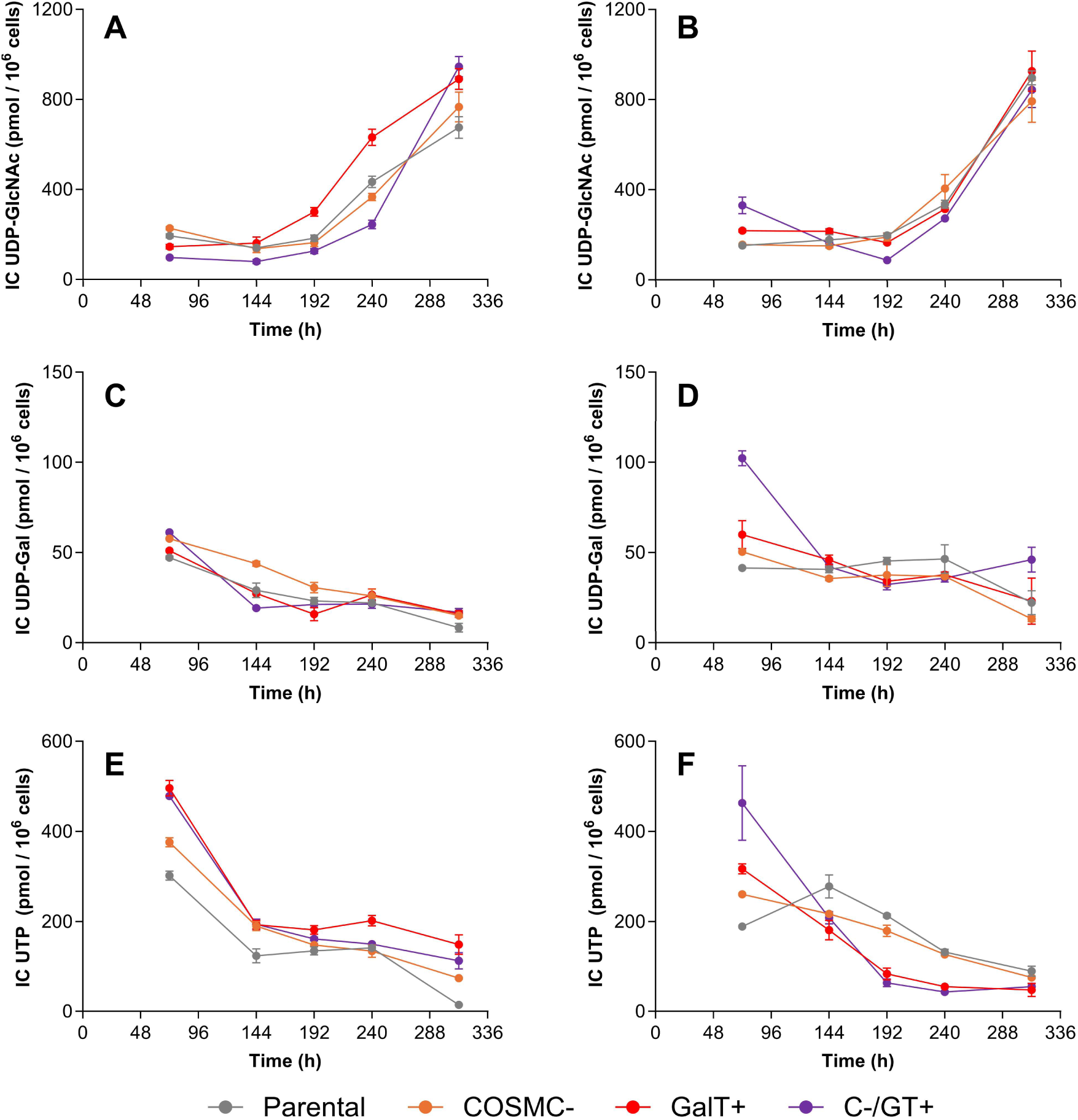
Intracellular nucleotide and nucleotide sugar donor profiles for batch cultures of DP12 and VRC01 glycovariant cells. Intracellular UDP-GlcNAc pool of DP12 (A) and VRC01 (B) glycovariants. Intracellular UDP-Gal pool of DP12 (C) and VRC01 (D) glycovariants. Intracellular pool of UTP in DP12 (E) and VRC01 glycovariant cells (F). Data for Parental (grey), COSMC- (orange), GalT+ (red), and C-/GT+ (purple) cells are shown. Datapoints are mean ± S.D. of N = 2 replicates.

### 3.7. Effects of glycoengineering on intracellular nucleotide and NSD pools

To determine how glycoengineering influences glycosylation metabolism, intracellular profiles of nucleotide and nucleotide sugar donor (NSD) pools in fed-batch cultures were determined using ion-pairing HPLC (Nakajima et al., 2010). Eight NSDs and 12 nucleotide phosphates were resolved with good linear quantification (Supplementary Figures 9-10 and Supplementary Table 6). The profiles of uridine diphosphate N-acetylglucosamine (UDP-GlcNAc), uridine diphosphate galactose (UDP-Gal), and uridine triphosphate (UTP) are presented in Figure 6.

In DP12 Parental, COSMC, and C-/GT+ cells, intracellular UDP-GlcNAc levels decrease slightly between 72 and 144 hours of culture to then accumulate until the end (Figure 6A). Early depletion may occur due to consumption towards production of cellular N-glycans, while subsequent accumulation may arise due to lower growth and, thereby, lower consumption towards cellular glycosylation. These trends are consistent with previously reported intracellular NSD profiles of CHO cells (Kochanowski et al., 2008). Interestingly, GalT+ cells have higher UDP-GlcNAc levels while C-/GT+ cells have the lowest (Figure 6A), suggesting that UDP-GlcNAc availability does not drive Man5 production by GalT+ cells (Figure 3A).

UDP-Gal profiles for DP12 glycovariants start with the highest levels at 72 hours of culture and, in general, decrease as culture progresses (Figure 6C). Parental, GalT+ and C-/GT+ cells have similar intracellular UDP-Gal profiles. In contrast, COSMC- cells present higher UDP-Gal between 72 and 240 hours of culture, which is consistent with the corresponding genetic modification. COSMC- cells lack UDP-Gal consumption towards β3-galactosylation of their O-linked glycans (Supplementary Figure 1) while simultaneously yielding Parental-like levels of mAb (Figure 3A) and cellular (Figure 5B) N-linked β4-galactosylation. The intracellular accumulation of UDP-Gal in DP12 COSMC- cells is likely due to the absence of the O-glycosylation sink and β4GalT-constrained N-linked galactosylation.

Intracellular UTP profiles for DP12 glycovariants start with high levels that decrease sharply until 144 hours of culture. Thereafter, UTP has a shallower decreasing trend to reach a minimum at the end of culture (Figure 6E). GalT+ and C-/GT+ variants present higher intracellular UTP pools, followed by COSMC- and Parental cells. Interestingly, higher intracellular UTP is observed in GalT+ and C-/GT+ cells, which were observed to achieve the lower peak VCD and IVCD (Table 2). It is possible that higher UTP demand occurs in Parental and COSMC- cells to sustain the RNA transcription required for higher growth.

The intracellular nucleotide and NSD pools for VRC01 glycovariants have similar trends to DP12 cells but present particular features. UDP-GlcNAc has an initially decreasing trend followed by accumulation, as in DP12 cells (Figure 6B). Parental, COSMC-, and GalT+ cells have similar profiles throughout. Although C-/GT+ cells have similar trends to the other glycovariants from 192 hours towards the end of culture, they start with a higher intracellular UDP-GlcNAc concentration and present a deeper drop until 192 hours of culture. The lower initial UDP-GlcNAc pool observed in GalT+ cells (compared with C-/GT+) may contribute to their higher production of Man5 mAb Fc glycoforms (Figure 3C).

The intracellular accumulation of UDP-Gal observed in DP12 COSMC- cells does not occur in VRC01 (Figure 6D). Although this NSD should accumulate in absence of the O-linked galactosylation sink, it may be that UDP-Gal biosynthesis is limited by UTP availability – VRC01 COSMC- cells present the second lowest initial UTP pool (Figure 6F). VRC01 C-/GT+ cells also present the highest initial level of UDP-Gal followed by the sharpest drop from 72 to 192 hours of culture (Figure 6D), suggesting that this NSD is rapidly consumed. This trend is consistent with the highest levels of mAb Fc (Figure 3D) and cellular (Figure 5D) N-linked galactosylation observed for these VRC01 glycovariants.

VRC01 glycovariants present slightly different trends in intracellular UTP pools than DP12 cells (Figure 6F). C-/GT+ and GalT+ VRC01 variants present the highest initial UTP pools that rapidly drop to the lowest levels from 192 hours of culture onwards. High UTP consumption also appears to correlate with cell growth in VRC01 glycovariants, where C-/GT+ and GalT+ cells show the highest consumption while presenting substantially higher peak VCD and IVCD than Parental and COSMC- variants (Table 2). Once again, high UTP consumption would be required for mRNA synthesis to sustain increased growth.

Overall, the intracellular nucleotide and NSD pools suggest that the knockout of COSMC has multiple positive effects that enhance mAb Fc β4-galactosyaltion while also reducing production of undesired Man5 glycans. The COSMC knockout reduces UDP-Gal demand towards cellular O-linked β3-galactosylation, freeing more of this substrate towards cellular and mAb N-linked β4-galactosylation. Reduced UDP-Gal demand leads to higher UTP availability towards UDP-GlcNAc biosynthesis, thus curbing Man5 production.

## 4. Discussion

A two-step glycoengineering strategy combining COSMC knockout with β4GalT1 overexpression (C-/GT+) consistently drives mAb N-linked galactosylation above 90% in both batch and fed-batch cultures while maintaining mAb titres comparable to or exceeding those of parental CHO cells. This level of hypergalactosylation is achieved through a streamlined workflow that uses lectin-aided FACS enrichment of COSMC knockout cells and balsticidin selection for β4GalT1 overexpression to deliver stable hypergalactosylating cell pools in less than six weeks. The obtained mAb Fc galactosylation levels advance what has been achievable through uridine/manganese/galactose (UMG) feeding (Grainger and James, 2013; Gramer et al., 2011; Pranomphon et al., 2025; Schäpertöns et al., 2026) or through equivalently simple cell engineering approaches (Raymond et al., 2015; Stach et al., 2019). Each engineering event alleviates one of the key bottlenecks of β4-galactosyaltion: the COSMC knockout eliminates O-linked β3-galactosylation and, thereby, frees UDP-Gal towards N-linked β4-galactosylation to alleviate metabolic bottlenecks, while β4GalT overexpression alleviates bottlenecks around cellular machinery availability.

The glycosylation profiles observed across Parental, COSMC-, and GalT+ glycovariants support the hypothesised mechanism of our dual C-/GT+ engineering strategy. β4GalT1 overexpression alone (GalT+) increases mAb Fc β4-galactosylation relative to Parental cells, which is consistent with previous reports where 70% (Stach et al., 2019) and 85% (Nguyen et al., 2021) mAb Fc β4-galactosylation is observed upon β4GalT1 overexpression and with the established role of this GalT isoform as the most specific for mAb Fc galactosylation (Bydlinski et al., 2018). However, GalT+ cells also exhibit a propensity to produce deleterious Man5 glycans with potential implications for mAb serum half-life (Goetze et al., 2011; Yu et al., 2012) and immunogenicity (Dasgupta et al., 2007; Delignat et al., 2020).

We propose that β4GalT1 overexpression – in the absence of COSMC knockout – creates competing metabolic pressures where elevated UDP-Gal consumption may deplete the intracellular UTP pool and, thereby, hinder UDP-GlcNAc biosynthesis (Pranomphon et al., 2025). We hypothesise that excessive Mn^2+^ sequestration by overexpressed β4GalT1 may reduce Mgat1 activity (Unligil et al., 2000), collectively diverting glycan processing towards Man5. In contrast, COSMC knockout alone eliminates UDP-Gal flux towards O-linked β3-galactosylation, which can account for up to 45% of total UDP-Gal consumption (del Val et al., 2016). Absence of active C1GalT, as achieved through knockout of COSMC (Yang et al., 2014), may alleviate Mn^2+^ sequestration (Gonzalez-Ramirez et al., 2022).

In conjunction, the COSMC knockout alleviates the UDP-Gal, UDP-GlcNAc, and Mn^2+^ constraints that promote Man5 accumulation but does not, on its own, provide sufficient β4GalT activity to drive hypergalactosylation. It is also important to note that the wild-type O-linked glycans of CHO cells are characteristically simple mono- or bi-sialylated Core 1 structures (Donini et al., 2025; North et al., 2010), which is consistent with decades of adaptation to suspension growth, during which reliance on complex cell surface O-glycans may have been reduced. In line with this, COSMC knockout, which ablates Core 1 galactosylation, had no detectable impact on cell growth, suggesting that more processed O-glycan structures are not necessary for CHO cell proliferation.

The C-/GT+ combination eliminates both bottlenecks simultaneously. By augmenting β4GalT1 activity while redirecting UDP-Gal, UTP, and Mn^2+^ away from O-glycan synthesis, the strategy resolves the enzymatic, nucleotide sugar donor, and cofactor bottlenecks that limit β4-galactosylation and promote Man5 production. The result is consistent hypergalactosylation above 90%, an outcome that, to our knowledge, has not been achieved at comparable mAb titres and with as few as two engineering events (Nguyen et al., 2021; Raymond et al., 2015; Stach et al., 2019). Notably, this consistency is demonstrated in pooled cell populations, suggesting that even greater uniformity would be expected following single-cell cloning.

### 4.1. Mechanistic Basis of Hypergalactosylation in C-/GT+ Cells

The glycosylation profiles across the four Parental, COSMC-, GalT+, and C-/GT+ glycoengineered variants reveal a coherent mechanistic interpretation of how enzyme availability, nucleotide and NSD supply, and cofactor distribution interact to govern mAb N-linked Fc β4-galactosyaltion (Figure 7). In parental cells, basal β4GalT1 expression is the primary constraint on N-linked β4-galactosylation. Because β4GalT activity is low, UDP-Gal consumption towards N-linked glycosylation is relatively modest, the intracellular UTP pool remains sufficient for UDP-GlcNAc synthesis via the hexosamine pathway, and Mn^2+^ – the obligate cofactor for C1GalT, Mgat1, and β4GalT1 (Boeggeman and Qasba, 2002; Gonzalez-Ramirez et al., 2022; Unligil et al., 2000) – is distributed across all three enzymes. In conjunction, these facets limit production of the deleterious Man5 glycan.

**Figure 7.**
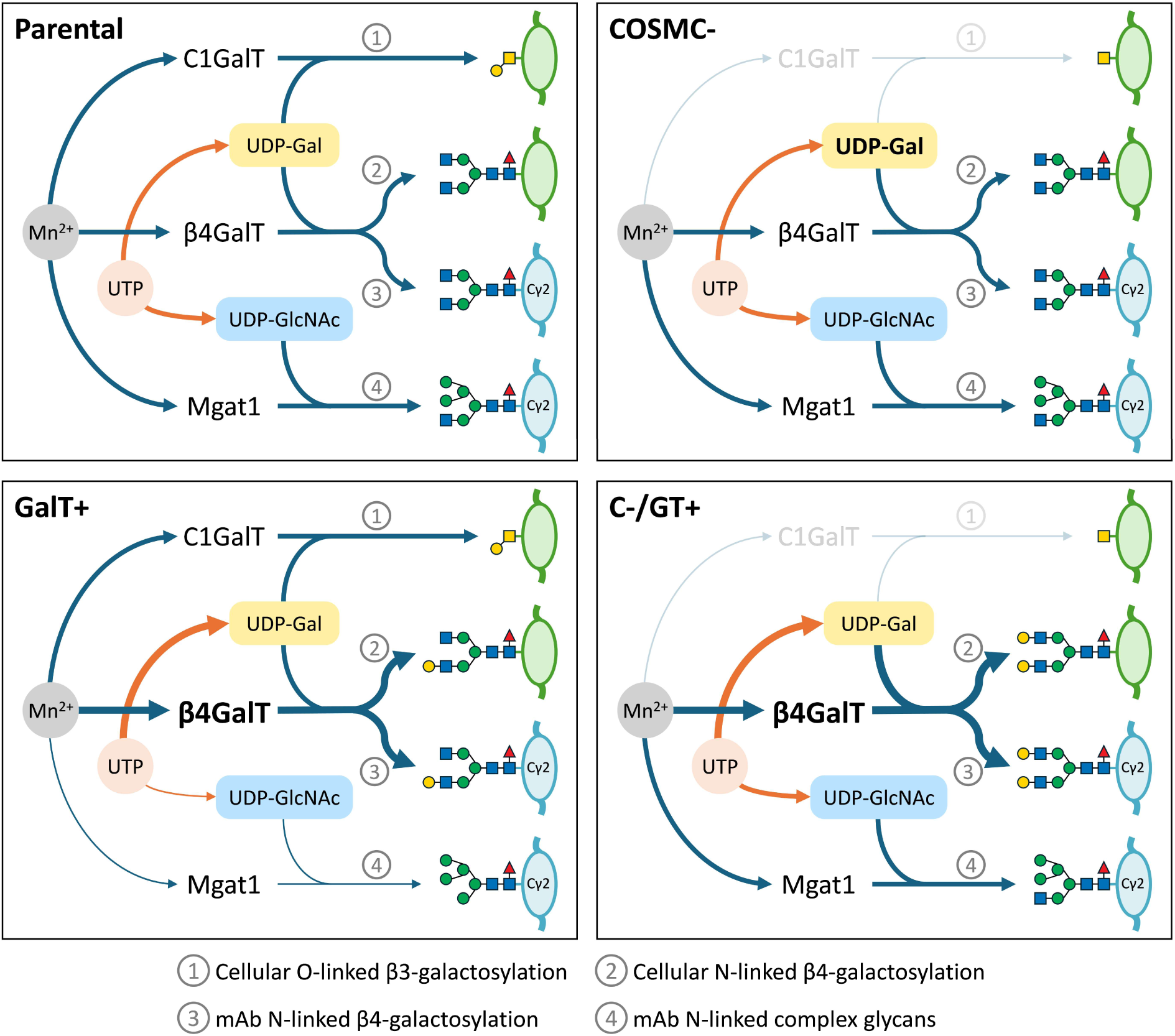
Proposed model for C-/GT+ hypergalactosylation. In parental cells, cellular and mAb N-linked β4-galactosylation are limited by β4GalT availability. At basal β4GalT expression levels, UDP-Gal consumption towards N-glycosylation is low, leaving UTP available for UDP-GlcNAc synthesis and, thereby, low levels of Man5 production. The Mn^2+^ cofactor is distributed among all enzymes. In COSMC- cells, C1GalT has no catalytic activity, yielding non-galactosylated Core 1 O-linked glycans. UDP-Gal accumulates due to absent O-linked β3-galactosylation, but because β4GalT availability remains limiting, N-linked β4-galactosylation is unaffected. With β4GalT overexpression (GalT+), cellular and mAb N-linked β4-galactosylation increases but becomes limited by UDP-Gal availability. Higher UDP-Gal consumption reduces UTP availability for UDP-GlcNAc synthesis, and β4GalT overexpression depletes Mn^2+^ for Mgat1 activity, in conjunction resulting in Man5 glycan production. Combining COSMC knockout with β4GalT overexpression (C-/GT+) redirects the UDP-Gal that is no longer consumed for cellular O-linked β3-galactosylation towards cellular and mAb N-linked β4-galactosylation. This pathway rewiring buffers excess UTP consumption to enable adequate UDP-GlcNAc production and, thus, reduce Man5 glycan production. Freed from C1GalT, Mn^2+^ becomes available for Mgat1, further reducing Man5 production. The C-/GT+ strategy simultaneously overcomes metabolic and cellular machinery bottlenecks to achieve mAb hypergalactosylation (>90%).

COSMC knockout disrupts Core 1 O-glycan β3-galactosylation by ablating correct folding of the C1GalT enzyme (Yang et al., 2014). Absence of C1GalT activity eliminates UDP-Gal consumption towards O-linked β3-galactosylation and results in intracellular accumulation of this NSD. However, because β4GalT1 availability remains the downstream bottleneck, the increased UDP-Gal pool cannot be redirected towards N-linked β4-galactosylation in cells that have only been modified with the COSMC knockout.

β4GalT1 overexpression (GalT+) substantially increases both cellular and mAb N-linked galactosylation by alleviating the cellular machinery bottleneck, consistent with prior reports (Nguyen et al., 2021; Raymond et al., 2015; Stach et al., 2019). However, GalT+ cells also exhibit elevated Man5 production, which we attribute to two inter-related consequences of increased β4GalT1 availability. First, the sharp increase in N-linked galactosylation demands proportionally greater UDP-Gal consumption. Because UDP-Gal is synthesised from UTP and glucose-1-phosphate through action of the UGP-2 (EC 2.7.7.9), GALT (EC 2.7.7.12), and GALE (EC 5.1.3.2) enzymes (Murrell et al., 2004), high sustained β4-galactosylation places an increased demand on the intracellular UTP pool. As UTP availability is shared with the hexosamine biosynthetic pathway, its depletion impairs UDP-GlcNAc synthesis, stalling N-glycan processing and promoting Man5 accumulation. To avoid shortfalls in UTP availability, uridine is often supplied, along with galactose and Mn^2+^, in metabolic galactoengineering strategies (Grainger and James, 2013; Gramer et al., 2011). Second, the elevated abundance of β4GalT1 may sequester a disproportionate share of intracellular Mn^2+^. Because Mn^2+^ is also an obligate cofactor for Mgat1 (Unligil et al., 2000) – the enzyme responsible for the committed step from Man5 to complex-type N-glycan processing (Stanley et al., 2022) – its relative depletion at the Mgat1 node would independently impair high-mannose conversion and further drive production of Man5. Together, UDP-GlcNAc depletion and enzyme competition for Mn^2+^, provide a consistent explanation for why, when overexpressed on its own, β4GalT yields undesired Man5 glycans despite achieving gains in mAb β4-galactosylation.

The C-/GT+ strategy resolves all three constraints in concert (Figure 7). COSMC knockout redirects UDP-Gal away from O-linked β3-galactosylation, providing a larger substrate pool to sustain the elevated galactosyltransferase activity conferred by β4GalT1 overexpression without proportionally increasing UTP demand. This pathway rewiring buffers excessive demand for UTP, preserving UDP-GlcNAc biosynthesis and enabling efficient N-glycan processing through the complex-type pathway. Simultaneously, absence of C1GalT activity frees Mn^2+^ for redistribution to Mgat1, supporting the processing activity required to prevent Man5 accumulation. The net result is a configuration in which enzymatic capacity, NSD availability, and cofactor distribution are co-optimised to achieve consistent mAb Fc hypergalactosylation above 90% while suppressing Man5 secretion.

### 4.2. Robustness and process resilience

Beyond the extent of mAb Fc β4-galactosylation achieved, a practically important feature of the C-/GT+ strategy is its consistency. Hypergalactosylation above 90% is maintained across batch and fed-batch culture formats in pooled cell populations, suggesting that the underlying metabolic rewiring confers a degree of intrinsic robustness to the glycosylation phenotype. This is significant because mAb N-glycosylation – particularly β4-galactosylation – is well known to be sensitive to culture process variables, including fluctuations in nutrient availability and their downstream effects on NSD biosynthesis (Fan et al., 2015; Naik et al., 2018; Sumit et al., 2019), the supply of trace metals such as Mn²⁺ (Graham et al., 2019), accumulation of metabolic by-products including ammonia (Synoground et al., 2021), and excursions in culture pH (Reddy et al., 2025). By expanding the intracellular UDP-Gal pool through O-glycan pathway diversion rather than relying on exogenous supplementation, our C-/GT+ cell engineering may be inherently less susceptible to variability than other glycoengineering approaches. Whether the C-/GT+ strategy confers complete resilience to Mn^2+^ limitation, ammonia accumulation, or pH excursions remains to be rigorously tested, but the mechanistic logic – reduced competition for shared NSDs, NSD precursors, and cofactors – suggests this is a testable and plausible hypothesis warranting future investigation.

It is also notable that the glycosylation consistency reported here is achieved in pooled cell populations rather than clonal isolates. Single-cell cloning would reduce remaining cell-to-cell heterogeneity in transgene copy number and expression level, likely yielding even greater batch-to-batch reproducibility. This positions our C-/GT+ platform as a strong foundation for both early-stage process development, where speed and simplicity are paramount, and for progression towards clonal cell lines suitable for late-stage manufacturing.

### 4.3. Functional and Therapeutic Implications

The ability to consistently drive mAb Fc galactosylation above 90% has direct implications for the effector function and pharmacokinetic profile of therapeutic antibodies. Terminal β4-linked galactose residues on the Fc N-glycan are required for complement-dependent cytotoxicity (CDC), where they facilitate C1q binding, mAb hexamerisation, and activation of the classical complement pathway (Peschke et al., 2017; van Osch et al., 2021). Fc β4-galactosylation also modulates antibody-dependent cellular cytotoxicity (ADCC) (Thomann et al., 2016; Zhang et al., 2020) and antibody-dependent cellular phagocytosis (ADCP) (Kuhns et al., 2020), although the magnitude of these effects varies with antibody subclass and the specific effector cell populations engaged. For therapeutic applications where CDC or enhanced effector function is desired, maximising and standardising β4-galactosylation through cell engineering offers a more reliable route to consistent biological activity.

Sialyltransferases require terminal β4-linked galactose residues as substrates (Nguyen et al., 2021; Stanley et al., 2022). Therefore, hypergalactosylation therefore expands the substrate available for sialylation, which has been associated with anti-inflammatory activity and extended serum half-life in certain IgG subclasses (Sneed et al., 2025; Vattepu et al., 2022). Whether the C-/GT+ platform can be further leveraged to drive hypersialylation – by combining it with sialyltransferase overexpression and/or CMP-sialic acid pathway engineering – represents a logical and attractive extension of the current work.

Conversely, the suppression of Man5 glycans in C-/GT+ cells relative to GalT+ cells is itself therapeutically meaningful. Elevated Man5 promotes rapid mAb clearance via mannose receptor-mediated uptake in the liver (Baumeister et al., 2026; Falck et al., 2021), reducing serum half-life, and has also been associated with increased immunogenic risk through lectin-mediated immune recognition (Dasgupta et al., 2007; Delignat et al., 2020). Avoiding Man5 accumulation while achieving hypergalactosylation is not merely a glycan quality attribute. It is a prerequisite for a therapeutically viable product, and it is a distinction that separates the C-/GT+ strategy from β4GalT overexpression approaches that ignore metabolic bottlenecks which lead to production of Man5 mAbs.

## 5. Conclusions and Outlook

Taken together, these results establish the C-/GT+ glycoengineering strategy as a simple, rapid, and highly effective approach to maximising mAb N-linked β4-galactosylation. By addressing enzymatic, metabolic, and co-factor constraints simultaneously through just two engineering events, and by exploiting efficient enrichment strategies that compress the cell glycoengineering timeline to under six weeks, the platform offers a practical and scalable route to hypergalactosylating CHO cell pools. The C-/GT+ approach also confers resilience to process-related excursions (e.g., nutrient and trace metal availability), addressing the β4-galactosylation variability often observed in wild-type mAbs. The mechanistic framework presented here, centred on redirecting UDP-Gal, UTP, and Mn^2+^ from competing glycosylation pathways to N-linked β4-galactosylation, provides a conceptual foundation for rational design of future glycoengineering strategies targeting other glycan motifs. Broader application of this framework, including integration with sialylation engineering or extension to other mAb formats, will be an important area for future work.

## Acknowledgments

IGA and IJV thank the Mexican Council for Science & Technology (CONACyT Grant No. 438330) and the Irish Research Council (GOIPG/2017/1049) for their financial support. The authors gratefully acknowledge funding from Research Ireland through the Solid-State Pharmaceutical Cluster – RI’s Research Centre for Pharmaceuticals (12/RC/2275_P2 SSPC). We also thank Prof Brian Glennon and Dr Mark Sheehan of the Applied Process Company for their kind assistance with the Cedex^®^ BioAnalyzer work. The authors thank the National Vaccine Center at the National Institutes of Health for the CHO VRC01 cell line.

## Declaration of generative AI and AI-assisted technologies in manuscript preparation

During preparation of this work, the authors used Anthropic PBC’s Claude Sonnet 4.6 to improve the text’s clarity and conciseness. After using the tool, the authors reviewed and edited the content as needed and take full responsibility for the content of the published article.

## Author contributions

IGA: investigation, methodology, data analysis, and writing – original draft. MG: investigation, methodology, and data analysis. AT: data analysis and writing – original draft. AB: resources, supervision, and data analysis. SC: investigation, methodology, and data analysis. JB: resources, supervision, funding acquisition, and writing – review and editing. IJV: conceptualisation, data analysis, funding acquisition, project administration, resources, supervision, and writing – review and editing.

