## Supplementary Information for "Computationally inspired glycoengineering to maximise mAb β4-galactosylation"

*Author to whom correspondence should be addressed.

### Supplementary Table 1. COSMC cloning oligos

| Oligo name | | 5'- Sequence - 3' | Size (bp) | Backbone |
|---|---|---|---|---|
| sgRNA COSMC | T | CACCGAATATGTGAGTGTGGATGG | 24 | pX458 |
| | B | AAACCCATCCACACTCACATATTC | 24 | |

T = Top; B = Bottom

### Supplementary Table 2. pUNO and pUNO1-hb4galt1sequencing primers

| Sequencing primer name | 5'- Sequence - 3' | Size (bp) | Backbone |
|---|---|---|---|
| pUNO-1.1_Fwd | TCCGCCGTCTAGGTAAGTTTAAAG | 24 | pUNO1 |
| pUNO-1.2_Fwd | ACTGCGTCTCTCCTCACAAG | 20 | pUNO1-hb4galt1 |

### Supplementary Table 3. Amplification primers for hβGalTI

| Primer name | 5'- Sequence - 3' | Size (bp) | $T_m$ (°C) |
|---|---|---|---|
| GalTI_Fwd | CTTCGGGAGCCGCTCCTG | 18 | 63.8 |
| GalTI_Rv | TCTTATCCGTGTACCAAAACGC | 22 | |

**Supplementary Table 3.** NSD Ion-pairing chromatography sequence.

| Time (min) | Buffer A (%) | Buffer B (%) | Comment |
|---|---|---|---|
| -0.02 | 100 | 0 | Equilibration |
| 0 | 100 | 0 | Injection |
| 13 | 100 | 0 | Isocratic elution |
| 35 | 23 | 77 | Gradient elution |
| 36 | 0 | 100 | Re-equilibration |
| 50 | 0 | 100 | Equilibration |

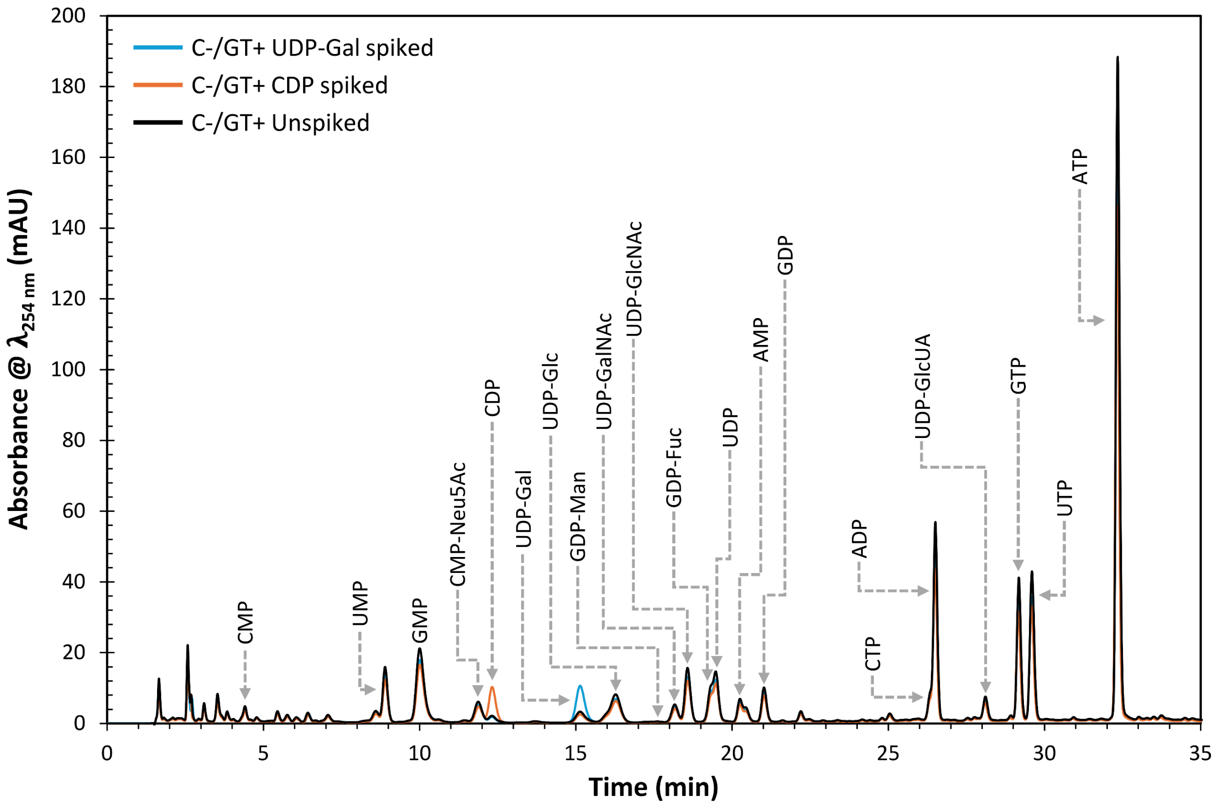


**Supplementary Figure 1. Sample chromatogram of intracellular nucleotide and nucleotide sugar donors (NSDs) extracted from DP12 C-/GT+ cells.**

### Supplementary Table 4. Standards for nucleotide and nucleotide sugar donor quantification

| Reagent | Abbreviation | Supplier | Cat. No. | Chemical formula | Molecular weight (g·mol$^{-1}$) |
|---|---|---|---|---|---|
| Adenosine 5′-monophosphate sodium salt | AMP | ACROS Organics | 440710010 | $C_{10}H_{13}N_5NaO_7P \cdot xH_2O$ | 369.21 |
| Cytidine 5′-monophosphate disodium salt | CMP | Sigma | C1006 | $C_9H_{12}N_3O_8PNa_2$ | 367.16 |
| Guanosine 5′-monophosphate disodium salt hydrate | GMP | Sigma | G8377 | $C_{10}H_{12}N_5Na_2O_8P \cdot xH_2O$ | 407.18 |
| Uridine 5′-monophosphate disodium salt | UMP | Sigma | U6375 | $C_9H_{11}N_2Na_2O_9P$ | 368.15 |
| Adenosine 5′-diphosphate sodium salt | ADP | Sigma | A2754 | $C_{10}H_{15}N_5O_{10}P_2$ | 427.20 |
| Cytidine-5'-diphosphate trisodium salt | CDP | Alfa Aesar | 15446339 | $C_9H_{21}N_3Na_3O_{11}P_2$ | 478.19 |
| Guanosine 5′-diphosphate sodium salt | GDP | Sigma | G7127 | $C_{10}H_{15}N_5O_{11}P_2$ | 443.2 |
| Uridine 5′-diphosphate disodium salt hydrate | UDP | Sigma | 94330 | $C_9H_{12}N_2Na_2O_{12}P_2 \cdot xH_2O$ | 448.12 |
| Adenosine 5′-triphosphate disodium salt hydrate* | ATP | ThermoFisher Scientific | R0441 | $C_{10}H_{16}N_5O_{13}P_3$ | 507.18 |
| Cytidine 5′-triphosphate disodium salt* | CTP | ThermoFisher Scientific | R0451 | $C_9H_{16}N_3O_{14}P_3$ | 483.16 |
| Guanosine 5′-triphosphate sodium salt hydrate* | GTP | ThermoFisher Scientific | R0461 | $C_{10}H_{16}N_5O_{14}P_3$ | 523.18 |
| Uridine 5′-triphosphate tris salt* | UTP | ThermoFisher Scientific | R0471 | $C_9H_{15}N_2O_{15}P_3$ | 484.14 |
| Uridine 5'-diphosphogalactose disodium | UDP-Gal | Sigma | U4500 | $C_{15}H_{22}N_2Na_2O_{17}P_2$ | 610.27 |
| Uridine 5′-diphosphoglucose disodium salt | UDP-Glc | Sigma | 94335 | $C_{15}H_{22}N_2Na_2O_{17}P_2$ | 610.27 |
| Uridine 5′-diphospho-N-acetylgalactosamine sodium salt | UDP-GalNAc | Sigma | U5252 | $C_{17}H_{25}N_3Na_2O_{17}P_2$ | 651.32 |
| Uridine 5′-diphospho-N-acetylglucosamine sodium salt | UDP-GlcNAc | Sigma | U4375 | $C_{17}H_{25}N_3O_{17}P_2Na_2$ | 651.32 |
| Uridine 5′-diphosphoglucuronic acid trisodium salt | UDP-GlcUA | Sigma | U6751 | $C_{15}H_{19}N_2Na_3O_{18}P_2$ | 646.23 |
| Cytidine-5′-monophospho-N-acetylneuraminic acid sodium salt | CMP-Neu5Ac | Sigma | C8271 | $C_{20}H_{31}N_4O_{16}P$ | 636.43 |
| Guanosine 5′-diphospho-D-mannose sodium salt | GDP-Man | Sigma | G5131 | $C_{16}H_{23}N_5O_{16}P_2Na_2$ | 649.30 |
| Guanosine 5'-diphospho-ß-L-fucose sodium salt | GDP-Fuc | Sigma | G4401 | $C_{16}H_{23}N_5Na_2O_{15}P_2$ | 633.31 |
| Guanosine 5′-diphosphoglucose sodium salt | GDP-Glc | Sigma | G7502 | $C_{16}H_{25}N_5O_{16}P_2$ | 605.34 |

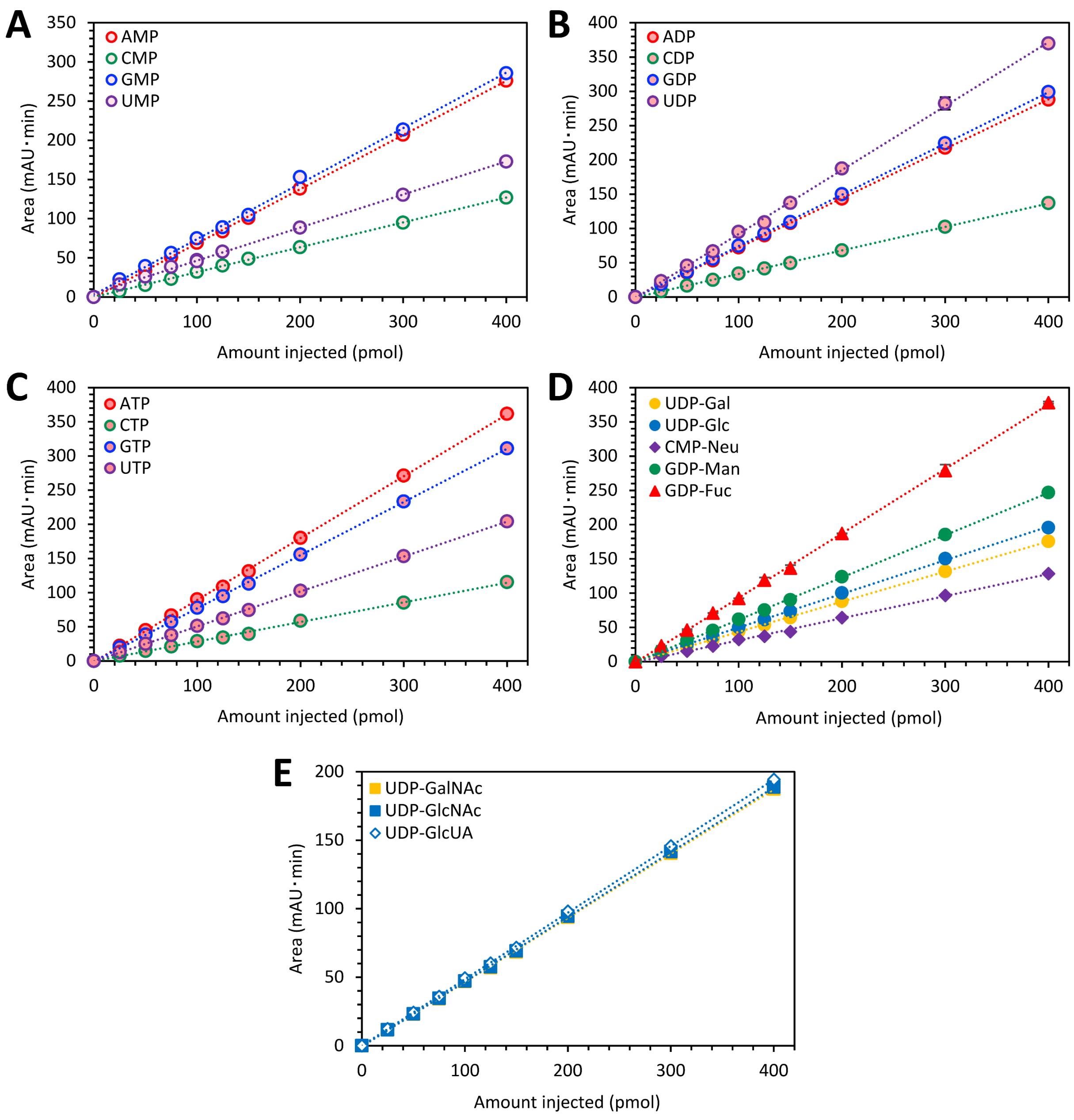


**Figure 2. Calibration curves for 12 nucleotide phosphates and eight nucleotide sugar donors (NSDs)**

**Supplementary Table 5.** NSD IP HPLC Reproducibility and Linearity

| Component | Reproducibility | | | Linearity | |
|---|---|---|---|---|---|
| | $t_{Ret}$ (min) | $t_{Ret}$ SD (min) | CV (%) | m ($mAU \cdot min \cdot pmol^{-1}$) | $R^2$ |
| AMP | 20.2 | 0.009 | 0.04 | 0.69 | 0.9998 |
| CMP | 4.8 | 0.006 | 0.13 | 0.32 | 0.9998 |
| GMP | 8.7 | 0.010 | 0.12 | 0.71 | 0.9984 |
| UMP | 7.1 | 0.016 | 0.23 | 0.42 | 0.9987 |
| ADP | 26.5 | 0.014 | 0.05 | 0.72 | 0.9998 |
| CDP | 12.3 | 0.006 | 0.05 | 0.34 | 0.9998 |
| GDP | 21.0 | 0.007 | 0.03 | 0.75 | 0.9999 |
| UDP | 19.4 | 0.013 | 0.07 | 0.93 | 0.9985 |
| ATP | 32.3 | 0.029 | 0.09 | 0.91 | 0.9997 |
| CTP | 26.3 | 0.037 | 0.14 | 0.29 | 0.9973 |
| GTP | 29.1 | 0.023 | 0.08 | 0.78 | 0.9998 |
| UTP | 29.5 | 0.026 | 0.09 | 0.51 | 0.9998 |
| UDP-Gal | 15.1 | 0.017 | 0.11 | 0.44 | 0.9999 |
| UDP-Glc | 16.2 | 0.018 | 0.11 | 0.49 | 0.9995 |
| UDP-GalNAc | 18.1 | 0.013 | 0.07 | 0.47 | 0.9999 |
| UDP-GlcNAc | 18.6 | 0.011 | 0.06 | 0.47 | 0.9999 |
| UDP-GlcUA | 28.0 | 0.022 | 0.08 | 0.49 | 0.9998 |
| CMP-Neu5Ac | 11.4 | 0.006 | 0.05 | 0.32 | 0.9984 |
| GDP-Man | 17.6 | 0.018 | 0.10 | 0.62 | 0.9998 |
| GDP-Fuc | 19.4 | 0.012 | 0.06 | 0.94 | 0.9989 |

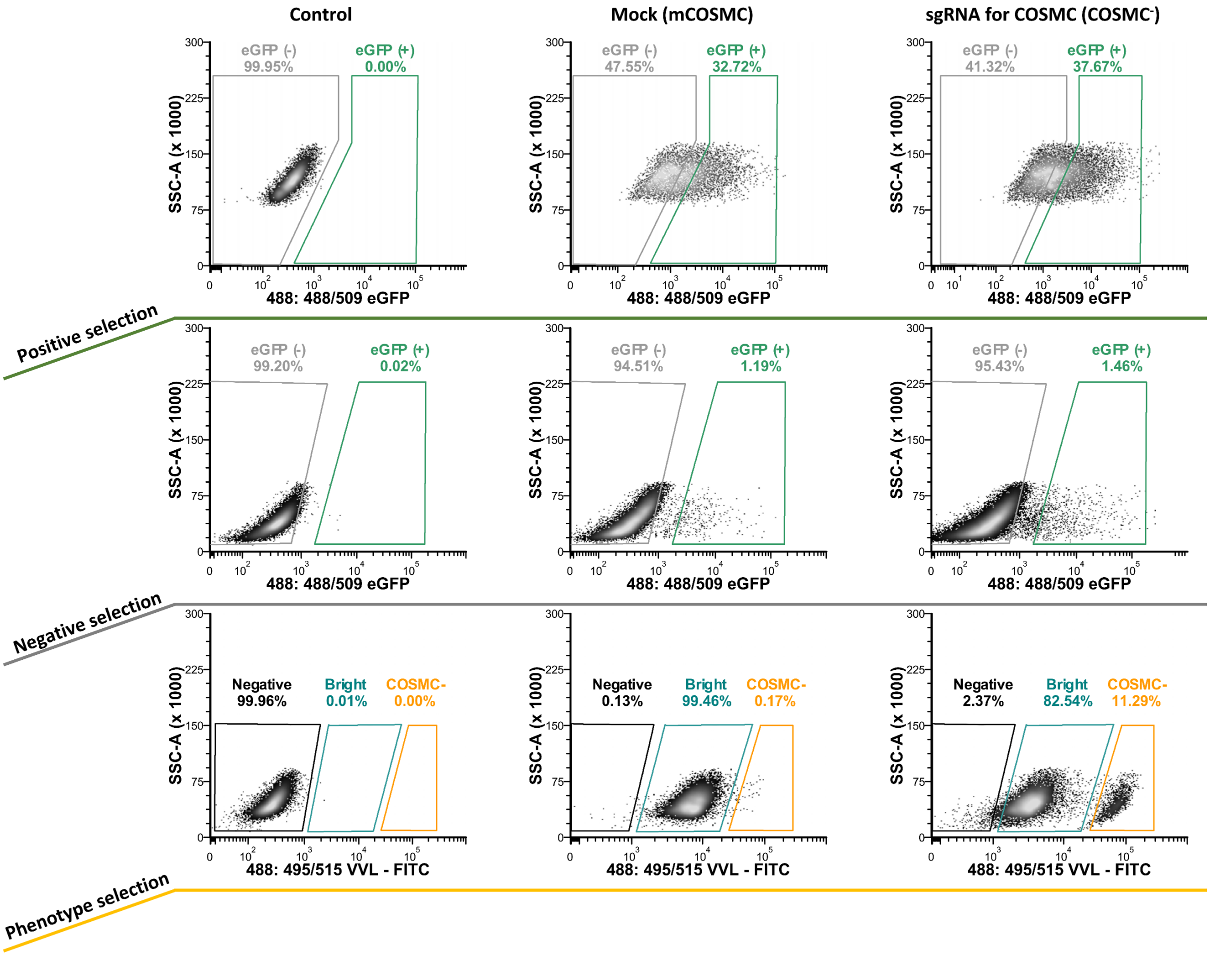


**Supplementary Figure 3.** Flow cytometry gate construction for identifying the *COSMC* knockout phenotype in CHO-DP12 cells.

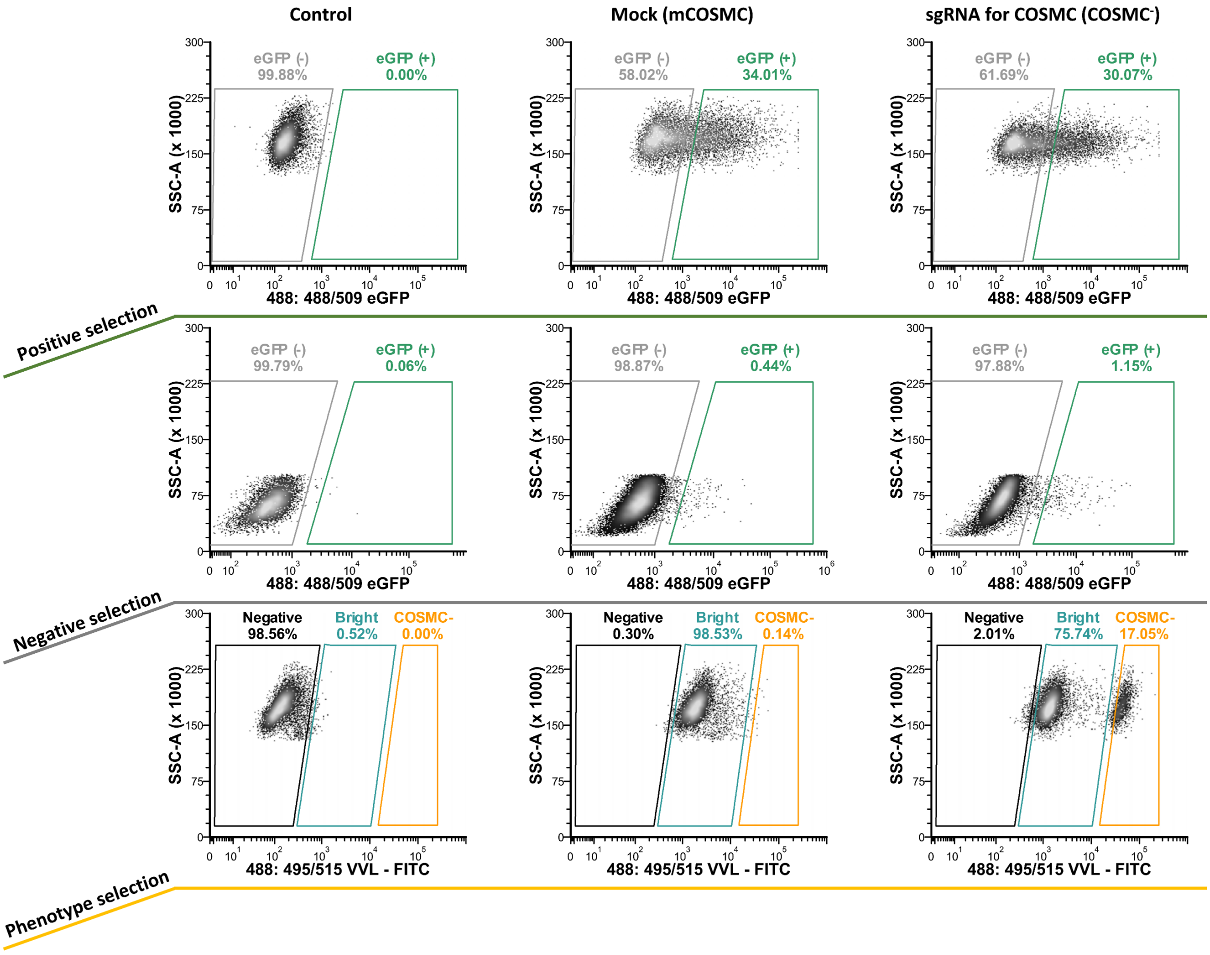


**Supplementary Figure 4.** Flow cytometry gate construction for identifying the *COSMC* knockout phenotype in CHO VRC01 cells.

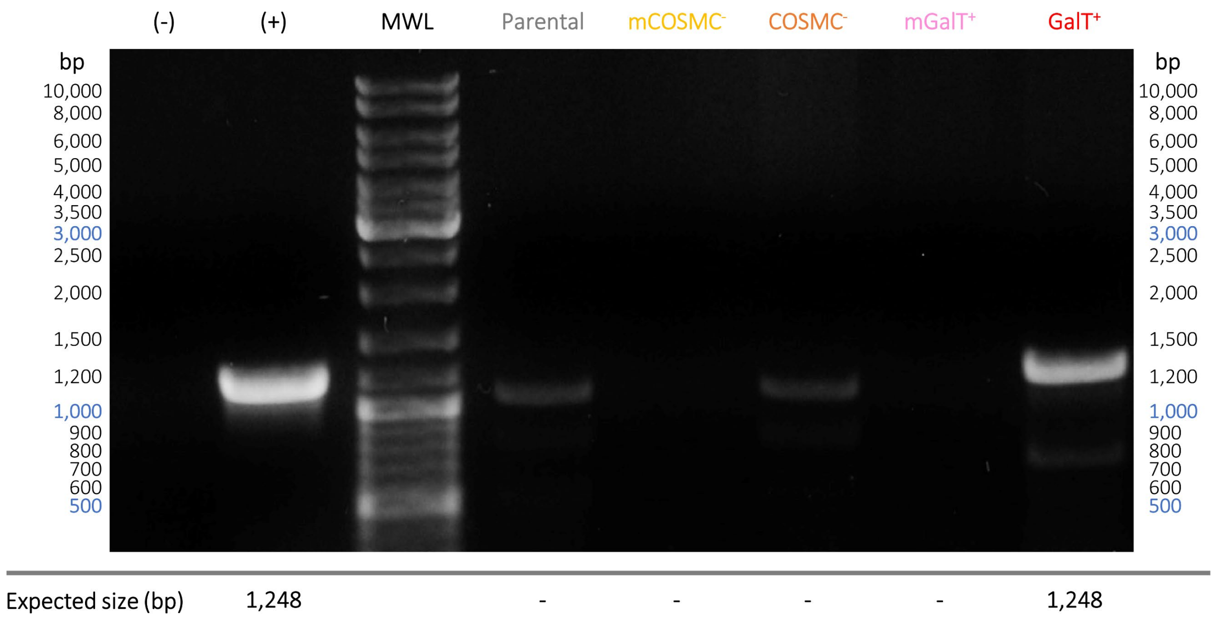


**Supplementary Figure 5. PCR-amplification of human galactosyltransferase I gene in CHO-DP12 pools**

(-) Negative control (PCR mix without primers), (+) Positive control (PCR mix using pUNO1-h4GalTI plasmid as DNA template).

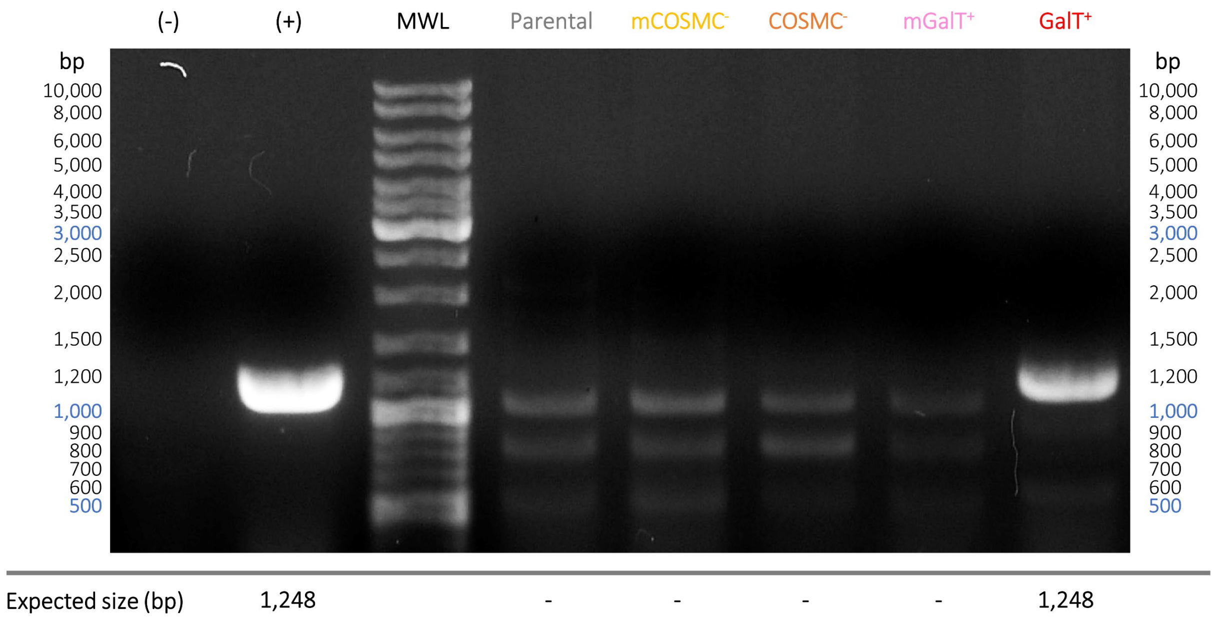


**Supplementary Figure 6. PCR-amplification of human galactosyltransferase I gene in CHO-VRC01 pools**

(-) Negative control (PCR mix without primers), (+) Positive control (PCR mix using pUNO1-h4GalTI plasmid as DNA template).

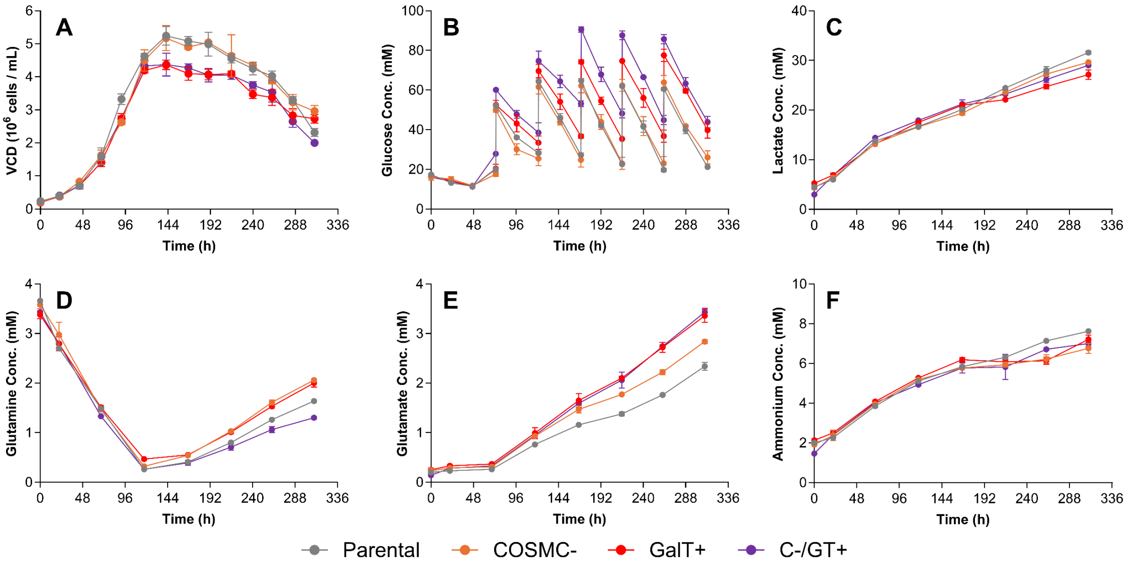


**Supplementary Figure 7. Metabolic profiles of CHO DP12 glycovariant cells cultured in fed batch mode**

Time profiles for VCD (A), Glucose (B), Lactate (C), Glutamine (D), Glutamate (E), and Ammonia (F) are shown. Grey corresponds to parental, orange to COSMC-, red to GalT+, and purple to C-/GT+ cells. The profiles correspond to mean of biological replicates (N=2) ± S.D.

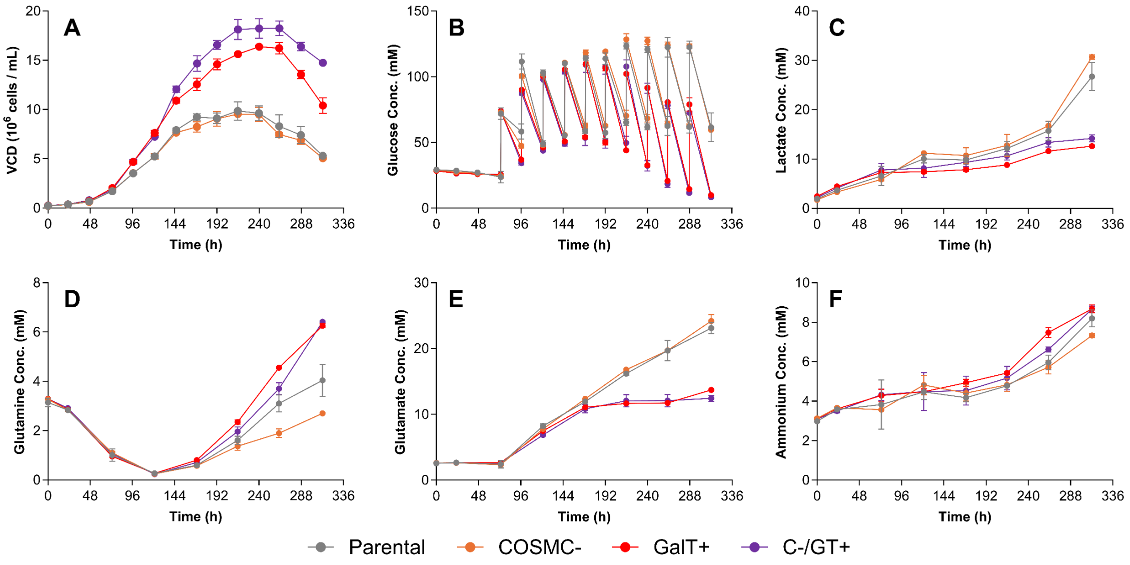


**Supplementary Figure 8. Metabolic profiles of CHO VRC01 glycovariant cells cultured in fed batch mode**

Time profiles for VCD (A), Glucose (B), Lactate (C), Glutamine (D), Glutamate (E), and Ammonia (F) are shown. Grey corresponds to parental, orange to COSMC-, red to GalT+, and purple to C-/GT+ cells. The profiles correspond to mean of biological replicates (N=2) ± S.D.

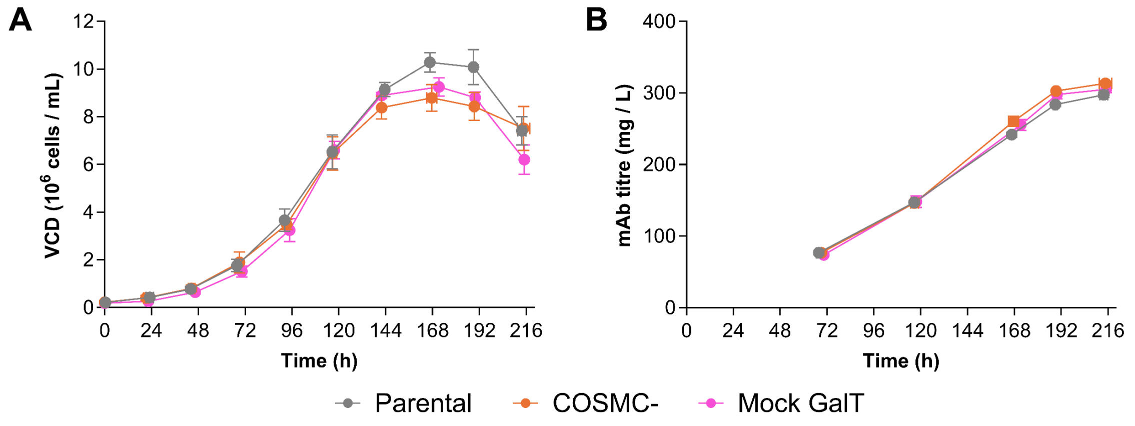


**Supplementary Figure 9. VCD and titre profiles for VRC01 mock GalT cells cultured in batch mode**

Time profiles for VCD (A) and mAb titre (B) are shown. Grey corresponds to parental, orange to COSMC-, and pink to mock GalT cells that have been transfected with pUNO plasmid devoid of the hβ4GalT gene and selected for blasticidin resistance. The profiles correspond to mean of biological replicates (N=2) ± S.D.

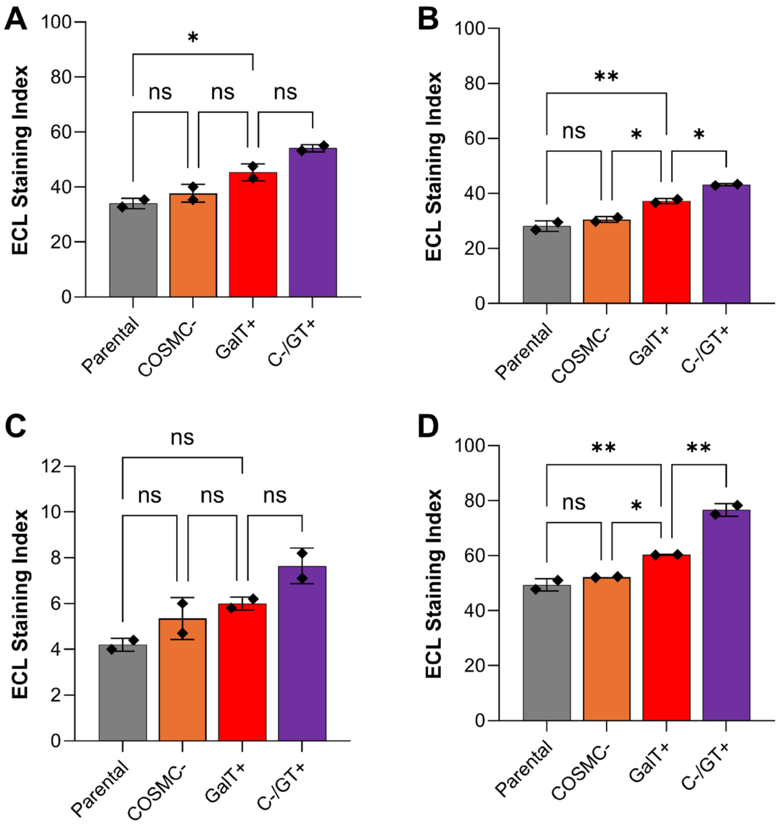


**Supplementary Figure 10. Non-normalised ECL Staining Index for DP12 and VRC01 glycovariant cells cultured in batch (A & C) and fed batch (B & D) mode**
